# Generalizing Bayesian phylogenetics to infer shared evolutionary events

**DOI:** 10.1101/2021.07.23.453597

**Authors:** Jamie R. Oaks, Perry L. Wood, Cameron D. Siler, Rafe M. Brown

## Abstract

Many processes of biological diversification can simultaneously affect multiple evolutionary lineages. Examples include multiple members of a gene family diverging when a region of a chromosome is duplicated, multiple viral strains diverging at a “super-spreading” event, and a geological event fragmenting whole communities of species. It is difficult to test for patterns of shared divergences predicted by such processes, because all phylogenetic methods assume that lineages diverge independently. We introduce a Bayesian phylogenetic approach to relax the assumption of independent, bifurcating divergences by expanding the space of topologies to include trees with shared and multifurcating divergences. This allows us to jointly infer phylogenetic relationships, divergence times, and patterns of divergences predicted by processes of diversification that affect multiple evolutionary lineages simultaneously or lead to more than two descendant lineages. Using simulations, we find the new method accurately infers shared and multifurcating divergence events when they occur, and performs as well as current phylogenetic methods when divergences are independent and bifurcating. We apply our new approach to genomic data from two genera of geckos from across the Philippines to test if past changes to the islands’ landscape caused bursts of speciation. Unlike our previous analyses restricted to only pairs of gecko populations, we find evidence for patterns of shared divergences. By generalizing the space of phylogenetic trees in a way that is independent from the likelihood model, our approach opens many avenues for future research into processes of diversification across the life sciences.

**Significance statement:** Phylogenetic models have long assumed that lineages diverge independently. Processes of diversification that are of interest in biogeography, epidemiology, and genome evolution, violate this assumption by affecting multiple evolutionary lineages. To relax the assumption of independent divergences and infer patterns of divergences predicted by such processes, we introduce a new way of conceptualizing, modeling, and inferring phylogenetic trees. We apply the new approach to genomic data from geckos distributed across the Philippines, and find support for patterns of shared divergences predicted by repeated fragmentation of the archipelago by interglacial rises in sea level.

## 1 Introduction

There are many processes of biological diversification that affect multiple evolutionary lineages, generating patterns of temporally clustered divergences across the tree of life. Understanding such processes of diversification has important implications across many fields and scales of biology. At the scale of genome evolution, the duplication of a chromosome segment harboring multiple members of a gene family causes multiple, simultaneous (or “shared”) divergences across the phylogenetic history of the gene family (Doyle and Egan, 2010; Jiao et al., 2011; Clark and Donoghue, 2017; Li et al., 2018). In epidemiology, when a pathogen is spread by multiple infected individuals at a social gathering, this will create shared divergences across the pathogen’s “transmission tree” (Pybus and Rambaut, 2009; Ypma et al., 2013; Klinkenberg et al., 2017). If one of these individuals infects two or more others, this will create a multifurcation (a lineage diverging into three or more descendants) in the transmission tree. At regional or global scales, when biogeographic processes fragment communities, this can cause shared divergences across multiple affected species (Hickerson et al., 2006; Leaché et al., 2007; Plouviez et al., 2009; Voje et al., 2009; Daza et al., 2010; Barber and Klicka, 2010). If the landscape is fragmented into three or more regions, this can also cause multifurcations (Hoelzer and Meinick, 1994). For example, the repeated fragmentation of the Philippines by interglacial rises in sea level since the late Pliocene (Haq et al., 1987; Rohling et al., 1998; Siddall et al., 2003; Miller et al., 2005; Spratt and Lisiecki, 2016) has been an important model to help explain remarkably high levels of microendemism and biodiversity across the archipelago (Inger, 1954; Heaney, 1985; Brown and Guttman, 2002; Evans et al., 2003; Heaney et al., 2005; Roberts, 2006; Linkem et al., 2010; Siler et al., 2010, 2011, 2012; Brown and Siler, 2014). This model predicts that recently diverged taxa across the islands should have (potentially multifurcating) divergence times clustered around the beginning of interglacial periods. We are limited in our ability to infer patterns of divergences predicted by such processes, because phylogenetic methods assume lineages diverge independently.

To formalize this assumption of independent divergences and develop ways to relax it, it is instructive to view phylogenetic inference as an exercise of statistical model selection where each topology is a separate model (Yang, 1994; Yang et al., 1995; Suchard et al., 2001). Current methods for estimating rooted phylogenies with *N* tips only consider tree models with *N* – 1 bifurcating divergences, and assume these divergences are independent, conditional on the topology (see Lewis et al., 2005, for multifurcations in unrooted trees). If, in the history leading to the tips we are studying, diversification processes affected multiple lineages simultaneously or caused them to diverge into more than two descendants, the true tree could have shared or multifurcating divergences. This would make current phylogenetic models with *N* – 1 independent divergence times over-parameterized, introducing unnecessary error (Figure 1). Even worse, with current methods, we lack an obvious way of using our data to test for patterns of shared or multifurcating divergences predicted by such processes.

**Figure 1.**
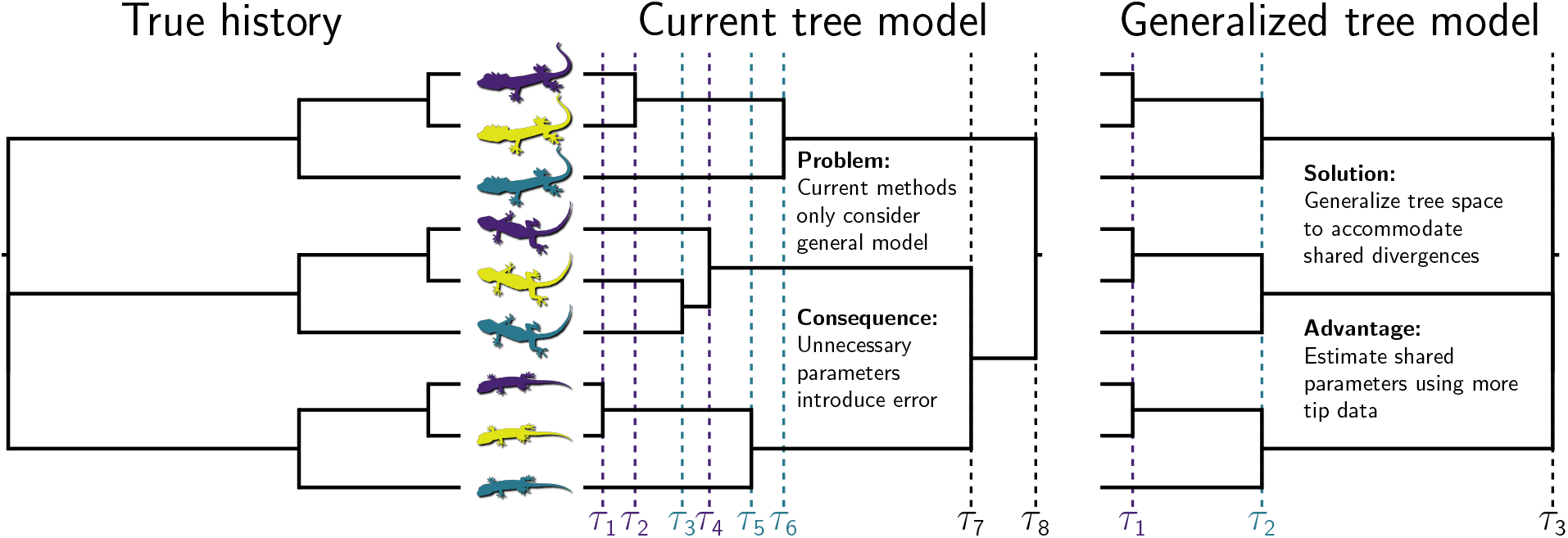
An example evolutionary history with shared divergences (left), and the benefits of the generalizing tree space under such conditions (right). Current methods are restricted to one class of tree models, where the tree is fully bifurcating and independent divergence-time parameters are estimated for all internal nodes (center). Figure made using Gram (Version 4.0.0; Foster, 2018) and the P4 phylogenetic toolkit (Version 1.4 5742542; Foster, 2004). Middle three lizard silhouettes from pixabay.com, and others from phylopic.org; all licensed under the Creative Commons (CC0) Public Domain Dedication.

We relax the assumption of independent, bifurcating divergences by introducing a Bayesian approach to generalizing the space of tree models to allow for shared and multifurcating divergences. In our approach, we view trees with *N* – 1 bifurcating divergences as only one class of tree models in a greater space of trees with anywhere from 1 to *N* – 1 potentially shared and/or multifurcating divergences (Figure S1). We introduce reversible-jump Markov chain Monte Carlo algorithms (Metropolis et al., 1953; Hastings, 1970; Green, 1995) to sample this generalized space of trees, allowing us to jointly infer evolutionary relationships, shared and multifurcating divergences, and divergence times. We couple these algorithms with a likelihood model for directly calculating the probability of biallelic characters given a population (or species) phylogeny, while analytically integrating over all possible gene trees under a coalescent model and all possible mutational histories under a finite-sites model of character evolution (Bryant et al., 2012; Oaks, 2019). Using simulations, we find the generalized tree model accurately infers shared and multifurcating divergences while maintaining a low rate of falsely inferring such divergences. To test for patterns of shared and multifurcating divergences predicted by repeated fragmentation of the Philippines by interglacial rises in sea level (Oaks et al., 2013; Brown et al., 2013; Oaks et al., 2019), we apply the generalized tree model to genomic data from two genera of geckos codistributed across the islands.

## 2 Results

### 2.1 Simulations on fixed trees

The generalized tree model (*M_G_*) sampled trees significantly closer (Robinson and Foulds, 1979; Kuhner and Felsenstein, 1994) to the true tree than an otherwise equivalent model that assumes independent, bifurcating divergences (*M_IB_*), when applied to 100 data sets simulated along the species tree in Figure 2A, each with 50,000 unlinked biallelic characters (Figure 2B). From these simulated data, the generalized model consistently inferred the correct shared and multifurcating divergences with high posterior probabilities (Figure 2C). Unlike the independent-bifurcating model, the generalized approach avoids strong support for nonexistent branches that spuriously split truly multifurcating nodes (Figure 2D). Under both models, analyzing only the variable characters causes a reduction in tree accuracy (Figure 2B), but yields similar posterior probabilities for shared and multifurcating divergences (Figure 2C).

**Figure 2.**
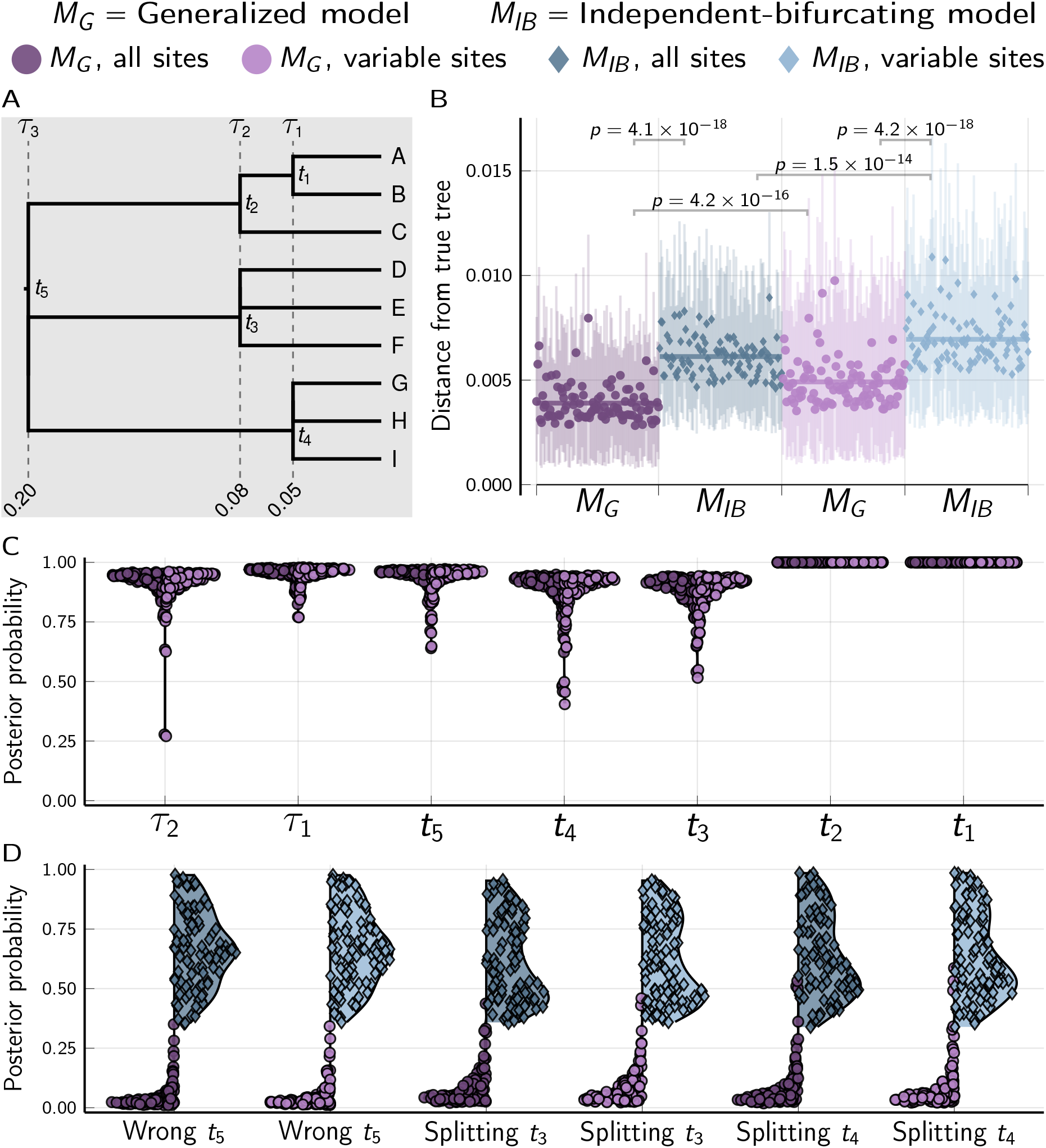
Results of analyses of 100 data sets, each with 50,000 biallelic characters simulated on the species tree shown in (A) with divergence times in units of expected substitutions per site. (B) The square root of the sum of squared differences in branch lengths between the true tree and each posterior tree sample (Kuhner and Felsenstein, 1994); the point and bars represent the posterior mean and equal-tailed 95% credible interval, respectively. P-values are shown for Wilcoxon signed-rank tests (Wilcoxon, 1945) comparing the paired differences in tree distances between methods. (C) Violin plots of the posterior probabilities of each node and shared divergence in the true tree across the 100 simulated data sets. (D) Violin plots of the most probable incorrect root node and most probable of the three incorrect splittings of the *t*_3_ and *t*_4_ multifurcations. For each simulation, the mutation-scaled effective population size (*N_e_μ*) was drawn from a gamma distribution (shape = 20, mean = 0.001) and shared across all the branches of the tree; this distribution was used as the prior in analyses. Tree plotted using Gram (Version 4.0.0, Commit 02286362; Foster, 2018) and the P4 phylogenetic toolkit (Version 1.4, Commit d9c8d1b1; Foster, 2004). Other plots created using the PGFPlotsX (Version 1.2.10, Commit 1adde3d0; Carlsson and Papp, 2021) backend of the Plots (Version 1.5.7, Commit f80ce6a2; Breloff, 2021) package in Julia (Version 1.5.4; Bezanson et al., 2017).

When applied to data sets of 50,000 characters simulated along a tree with independent, bifurcating divergences (Figure 3A), both the *M_G_* and *M_IB_* models consistently inferred the correct topology with strong support (Figure 2B), and the *M_G_* method did not support incorrect shared or multifurcating divergences (Figure 3C). This was true whether all the characters or only the variable characters were analyzed (Figure 3B&C). Looking at the distances (Robinson and Foulds, 1979; Kuhner and Felsenstein, 1994) between the trees from the posterior samples and the true tree, there is no difference between the *M_G_* and *M_IB_* models when the true tree has only independent, bifurcating divergences (Figure 3D). For both models, using all the characters yields posterior samples of more accurate trees than only analyzing variable characters (Figure 3D).

**Figure 3.**
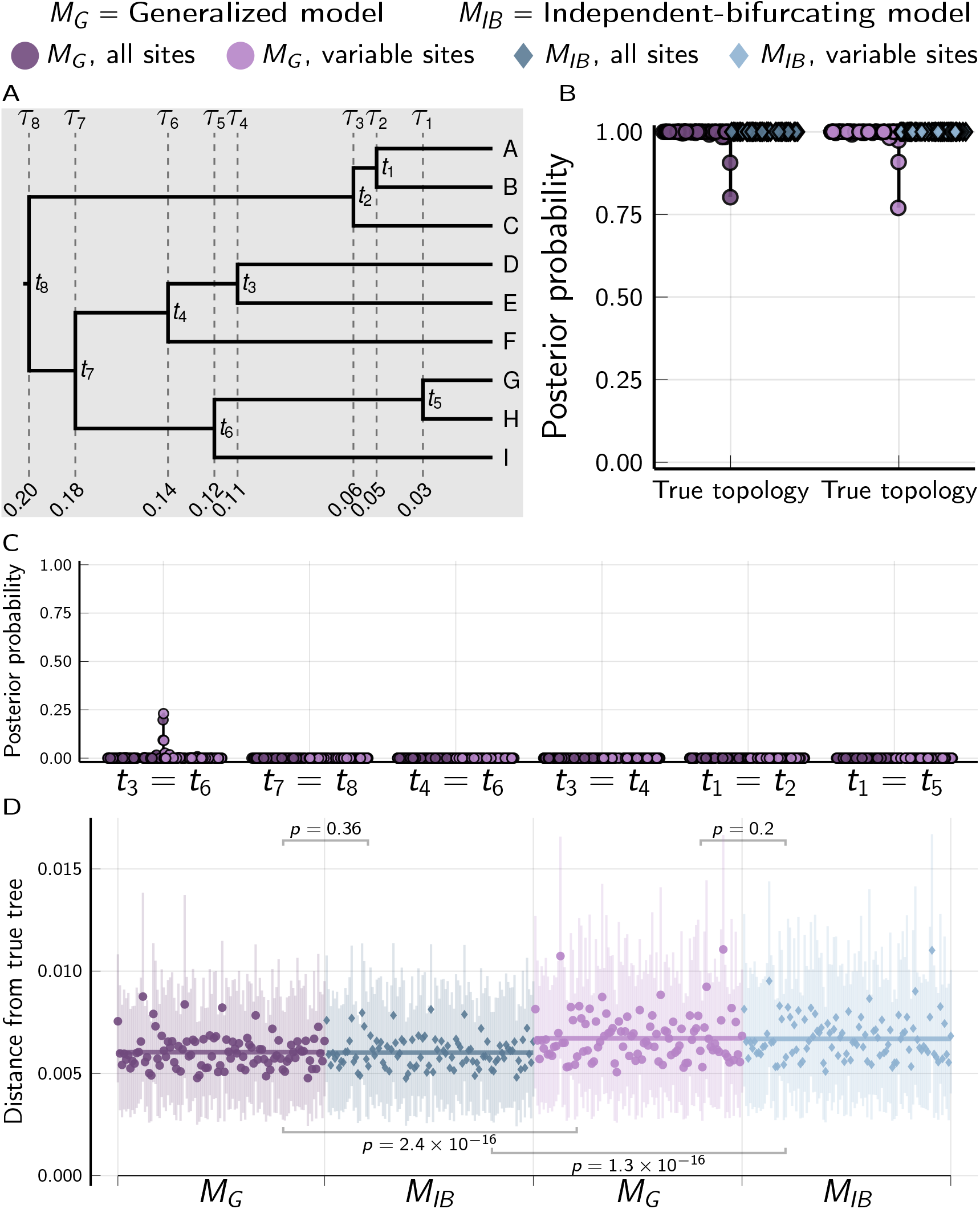
Results of analyses of 100 data sets, each with 50,000 biallelic characters simulated on the species tree shown in (A) with divergence times in units of expected substitutions per site. (B) The posterior probability of the true topology. (C) The posterior probability of incorrectly shared or multifurcating nodes. (D) The square root of the sum of squared differences in branch lengths between the true tree and each posterior tree sample (Kuhner and Felsenstein, 1994); the point and bars represent the posterior mean and equal-tailed 95% credible interval, respectively. P-values are shown for Wilcoxon signed-rank tests (Wilcoxon, 1945) comparing the paired differences in tree distances between methods. For each simulation, the mutation-scaled effective population size (*N_e_μ*) was drawn from a gamma distribution (shape = 20, mean = 0.001) and shared across all the branches of the tree; this distribution was used as the prior in analyses. Tree plotted using Gram (Version 4.0.0, Commit 02286362; Foster, 2018) and the P4 phylogenetic toolkit (Version 1.4, Commit d9c8d1b1; Foster, 2004). Other plots created using the PGFPlotsX (Version 1.2.10, Commit 1adde3d0; Carlsson and Papp, 2021) backend of the Plots (Version 1.5.7, Commit f80ce6a2; Breloff, 2021) package in Julia (Version 1.5.4; Bezanson et al., 2017).

### 2.2 Simulations on random trees

When we simulated 100 data sets (each with nine species and 50,000 characters) where the true tree and divergence times were randomly drawn from the generalized tree distribution (*M_G_*), we again found that the *M_G_* performs better than the *M_IB_* at inferring the correct tree and divergence times (Figure 4A), and generally recovers true shared and multifurcating divergences with moderate to strong support (Figure 4B&C). When the tree and divergence times were randomly drawn from an independent, bifurcating tree model (*M_IB_*), the generalized model performs similarly to the true model (Figure S2).

**Figure 4.**
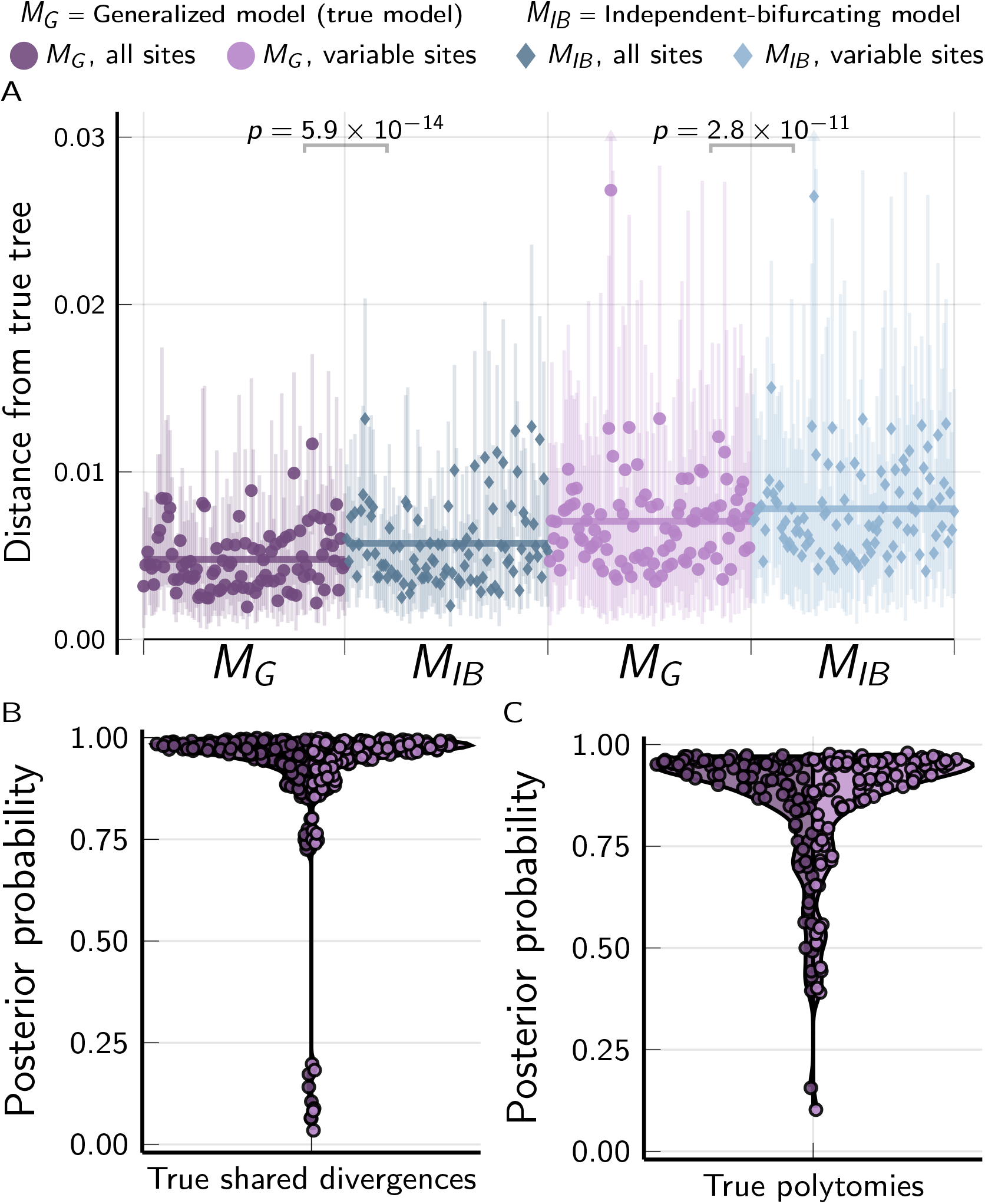
The performance of the *M_G_* and *M_IB_* tree models when applied to 100 data sets, each with 50,000 biallelic characters simulated on species trees randomly drawn from the *M_G_* tree distribution. (A) The square root of the sum of squared differences in branch lengths between the true tree and each posterior tree sample (Kuhner and Felsenstein, 1994); the point and bars represent the posterior mean and equal-tailed 95% credible interval, respectively. P-values are shown for Wilcoxon signed-rank tests (Wilcoxon, 1945) comparing the paired differences in tree distances between methods. Violin plots show posterior probabilities of all true (B) shared divergences and (C) multifurcating nodes across all simulated trees. For each simulation, the mutation-scaled effective population size (*N_e_μ*) was drawn from a gamma distribution (shape = 20, mean = 0.001) and shared across all the branches of the tree; this distribution was used as the prior in analyses. Plots created using the PGFPlotsX (Version 1.2.10, Commit 1adde3d0; Carlsson and Papp, 2021) backend of the Plots (Version 1.5.7, Commit f80ce6a2; Breloff, 2021) package in Julia (Version 1.5.4; Bezanson et al., 2017).

Both the *M_G_* and *M_IB_* models accurately and precisely estimate the age of the root, tree length, and effective population size from the data sets simulated on random *M_G_* and *M_IB_* trees (Top two rows of Figures S3, S4, and S5, respectively). Accuracy is similar with and without constant characters, but precision is higher when including constant characters.

### 2.3 The rate of falsely inferring shared divergences

To quantify the rate at which `phycoeval` incorrectly infers shared and/or multifurcating divergences, we used the results from the *M_G_* analyses of the data sets simulated on random trees from the *M_G_* and *M_IB_* models. From the posterior sample of each analysis, we used `sumphycoeval` to calculate the proportion of samples that contained incorrectly merged neighboring divergence times. To do this, we merged all possible neighboring divergence times from the true tree, each of which creates a shared divergence or multifurcation, and counted how many posterior samples contained each divergence scenario. We found that `phycoeval` had a low false-positive rate for the simulated data; less than 1% (Figure 5A & S6) and 5% (Figure 5D & S7) of incorrectly merged divergence times had an approximate posterior probability greater than 0.5 when analyzing data simulated on trees sampled from the *M_G_* and *M_IB_* models, respectively. In all cases with moderate to strong support for falsely merged divergences, the difference in time between the merged divergences was small (< 0.005 expected substitutions per site; Figure 5B&E). There was no correlation between support for incorrectly merged divergences and their age (Figure 5C&F; the p-value for a t-test that Pearson’s correlation coefficient = 0 using all points with posterior probability > 0 was 0.11 and 0.25 for results from data simulated under *M_G_* and *M_IB_*, respectively).

**Figure 5.**
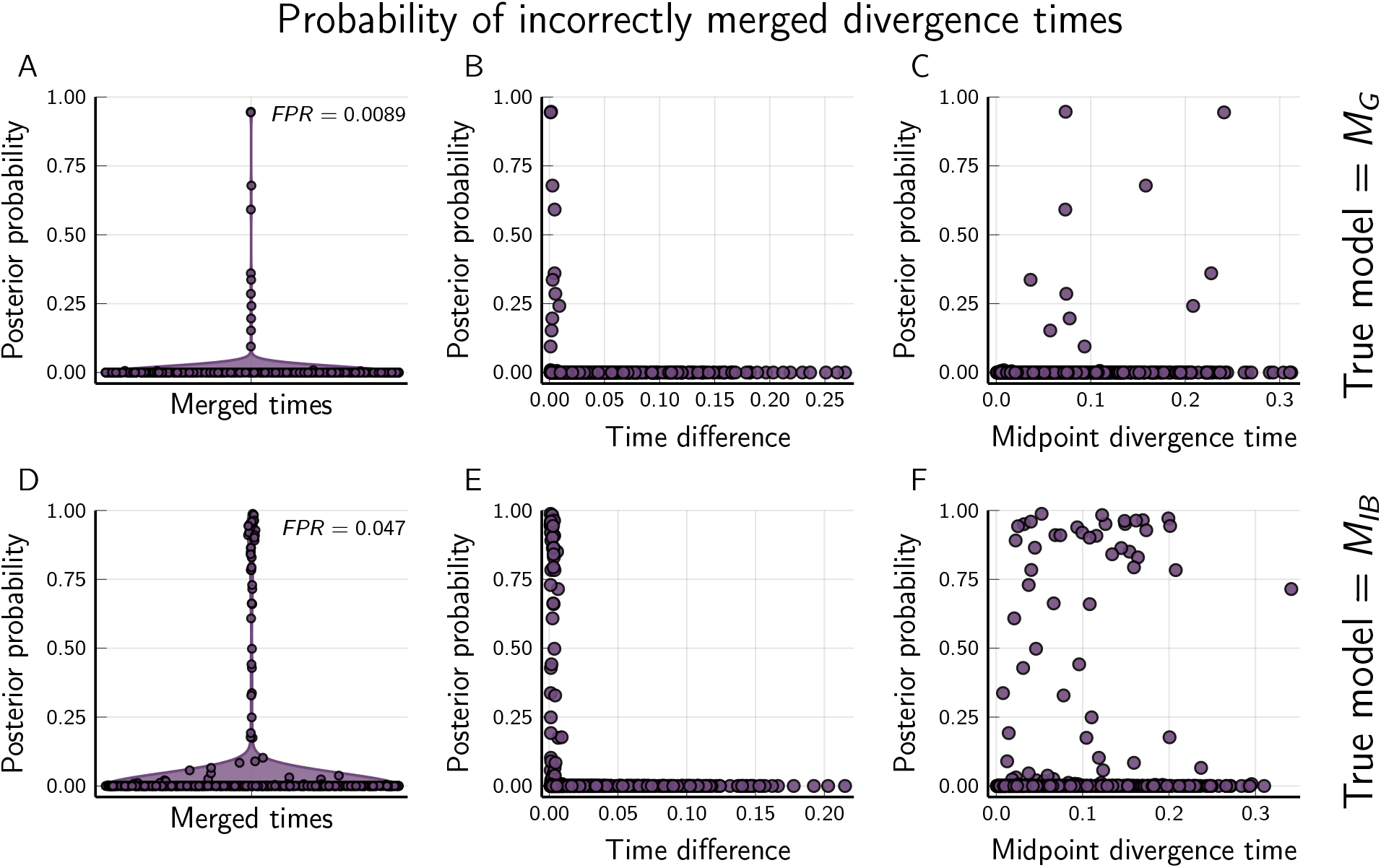
The *M_G_* tree model has (A & D) a low false positive rate (FPR; the proportion of incorrectly merged divergence times with a posterior probability > 0.5) when applied to data simulated on trees drawn from the (A–C) *M_G_* and (D–F) *M_IB_* models. Support for incorrectly merged divergence times is high only when the difference between the times is small (B & E), and is not correlated with the age of the merged nodes; p-value = (C) 0.11 and (F) 0.25 for a t-test that Pearson’s correlation coefficient = 0 using all points with posterior probability > 0. Time units are expected substitutions per site. Plots created using the PGFPlotsX (Version 1.2.10, Commit 1adde3d0; Carlsson and Papp, 2021) backend of the Plots (Version 1.5.7, Commit f80ce6a2; Breloff, 2021) package in Julia (Version 1.5.4; Bezanson et al., 2017).

### 2.4 Convergence and mixing of MCMC chains

For all analyses of simulated data, the root age, tree length, and effective population size had a potential-scale reduction factor (PSRF; the square root of Equation 1.1 in Brooks and Gelman, 1998) less than 1.2 and effective sample size (ESS; Gong and Flegal, 2016) greater than 200. The average standard deviation of split frequencies (ASDSF) among the four MCMC chains was less than 0.017 for all analyses and less than 0.01 for most (Figure S8).

Convergence and mixing was better under *M_G_* than *M_IB_* when applied to data sets simulated on trees with shared or multifurcating divergences (left column of Figure S8). When applied to data sets simulated with no shared or multifurcating divergences, MCMC performance was similar between *M_G_* and *M_IB_* (right column of Figure S8).

The MCMC settings used for *M_G_* and *M_IB_* are identical except for the reversible-jump moves that add or remove divergence-time parameters are turned off under the latter model. Under the *M_IB_* model, the tree topology is updated by several MCMC moves (see Sections 4.5 and 4.6.1 of the Supporting Information), which performed well when divergences are independent and bifurcating (Figure 3 and Figure S8). To further probe the improved MCMC behavior of MG in the face of shared and multifurcating divergences, we re-ran the analyses under the *M_IB_* model on the 100 data sets simulated along the tree in Figure 2A with more favorable MCMC settings. In these *M_IB_* reanalyses, we ran the MCMC chains twice as long, sampled them half as frequently, and started them with the correct tree. The results were nearly identical to the original MCMC chains under the *M_IB_* model (Figure S9), suggesting the improved mixing under MG was not simply do to insufficient MCMC sampling effort under the *M_IB_* model.

### 2.5 Simulations of linked characters

The multi-species coalescent likelihood we have coupled with our generalized tree model assumes each biallelic character is unlinked (i.e., each character evolved along a gene tree that was independent of other characters, conditional on the species tree; Bryant et al., 2012; Oaks, 2019). However, each locus comprising the gecko data sets we analyzed (see below) consists of approximately 90 contiguous nucleotides. To assess whether linked sites might bias our results, we repeated the simulations above, but with 500 loci, each with 100 linked characters. When all characters (variable and constant) are analyzed, results from the data sets simulated with linked characters are very similar to results from unlinked characters above (Figures S3–5 and 7 and S10–13). When all but one variable character per locus is discarded to avoid violating the assumption of unlinked characters, performance is greatly reduced due to the large loss of data (Figures S3–5 and 7 and S10–13). These results suggest the model is robust to linked characters and it is better to analyze all sites from multi-locus data sets, rather than reduce them to only one SNP per locus.

### 2.6 Testing for shared divergences in Philippine gekkonids predicted by glacial cycles

If the repeated fragmentation of the Philippines by interglacial rises in sea level generated pulses of speciation, taxa distributed across the archipelago should have divergence times clustered around the beginning of interglacial periods. We tested this prediction by applying our generalized tree model to RADseq data from species of *Cyrtodactylus* and *Gekko* collected from 27 and 26 locations across the islands, respectively (Tables S1 & S2). We analyzed each genus separately, because the rate of mutation differs between the genera, and `phycoeval` currently assumes a strict clock (though this is not required by the generalized tree model).

The maximum *a posteriori* (MAP) trees for both genera had 16 divergence times and weak to moderate support for five shared divergences (Figure 6; see Figures S14 & S15 and Table S3 for more details about shared divergences). The MAP tree of *Cyrtodactylus* and *Gekko* had three and two multifurcations, respectively. For both genera, two of the shared divergences involved three nodes, and of the remaining three that involved two nodes, one involved a trichotomous node (three descending lineages). There were no other strongly supported shared divergences that were not included in the MAP trees of either genera. Most of the shared and multifurcating divergences occurred after the late Pliocene (Figure 6 and Table S3), based on re-scaling the branch lengths of the posterior sample of trees from expected substitutions per site to millions of years using secondary calibrations (see methods).

**Figure 6.**
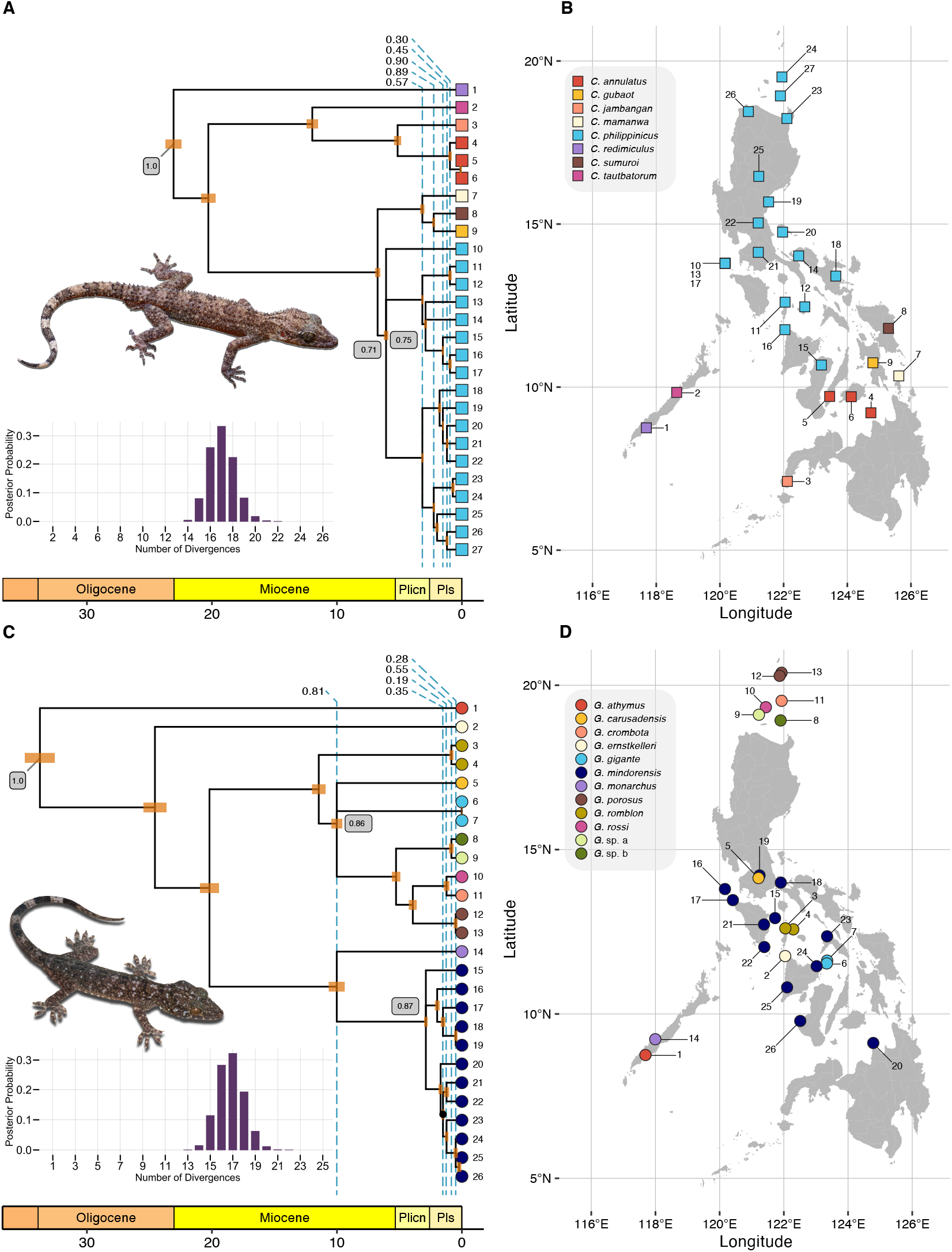
A summary of the generalized trees inferred from the (A–B) *Cyrtodactylus* and (C–D) *Gekko* RADseq data sets. The maximum a *posteriori* (MAP) tree is shown for both genera along with the approximate posterior probabilities of the number of divergences. Shared divergences in MAP trees indicated by dashed lines, with approximate posterior probabilities shown along the top. All clades (splits) had approximate posterior probabilities (PP) greater than 0.95 except for one indicated with a dot (PP = 0.89) within *G. mindorensis* (C). Approximate posterior probabilities of nodes shown in grey boxes for the root and multifurcating nodes. To illustrate timescale, branch lengths of posterior samples of trees were rescaled from expected substitutions per site to millions of years using secondary calibrations (see methods). Top photo of *Cyrtodactylus* sp. by CDS; bottom photo of *Gekko* sp. by Jason Fernandez & RMB. Created using ggplot2 (v3.3.5; Wickham, 2016), ggtree (v3.1.0; Yu et al., 2017), treeio (v1.17.0; Wang et al., 2019), deeptime (v0.0.6; Gearty, 2021), cowplot (v1.1.1; Wilke, 2020), and ggrepel (v0.9.1; Slowikowski, 2020). Links to nexus-formatted annotated trees: *[Cyrtodactylus](https://raw.githubusercontent.com/phyletica/phycoeval-experiments/master/gekkonid-output/map-tree-cyrt-nopoly.nex)* & *[Gekko](https://raw.githubusercontent.com/phyletica/phycoeval-experiments/master/gekkonid-output/map-tree-gekko-nopoly.nex)*.

For both genera, the number of divergence times with the highest approximate posterior probability (0.33 for *Cyrtodactylus* and 0.32 for *Gekko*) was 17, and the 95% credible interval spanned 15–19 divergences (Figure 6). No trees with more than 22 divergence times were sampled for either genera, making the approximate posterior probability of 23 or more divergences less than 2.9×10^−5^ for both genera. The average standard deviation of split frequencies (0.0027 for *Cyrtodactylus* and 0.0009 for *Gekko*) and other statistics were consistent with the MCMC chains converging and mixing well (Table S4).

## 3 Discussion

To relax the assumption that all processes of biological diversification affect evolutionary lineages independently, we introduced a generalized Bayesian phylogenetic approach to inferring phylogenies with shared and multifurcating divergences. Using simulations we found this approach can accurately infer shared and multifurcating divergences from moderately sized data sets, while maintaining a low rate of incorrectly inferring such patterns of divergence. When we used the generalized approach to infer the evolutionary histories of two genera of gekkonid lizards across the Philippines, we found strong support against tree models assumed by current phylogenetic methods. The posterior probability of all trees with *N* – 1 independent, bifurcating divergences was less than 2.9×10^−5^ for both genera, suggesting that trees with shared and multifurcating divergences better explain the gekkonid sequence data. It will be interesting to see if such improvement in model fit is common as the generalized tree distribution is applied to more systems, regardless of the biological processes responsible (if any).

Despite greatly expanding the number of possible topologies, we saw better MCMC behavior under the *M_G_* model (Figure S8), even when the *M_IB_* chains were started with the true tree and run twice as long (Figure S9). This could be due to the generalized tree distribution providing more ways to traverse tree space. For example, when a posterior distribution restricted to trees with independent bifurcating divergences has multiple “peaks” associated with different topologies, the generalized distribution includes tree models that are special cases of these topologies. Explicitly including these “intermediate” trees could make the posterior less rugged and allow MCMC chains to more easily traverse tree space.

By accommodating multifurcations, our generalized tree approach helped avoid the “star-tree paradox,” where arbitrary resolutions of a true polytomy can be strongly supported (Figure 2D; Suzuki et al., 2002; Lewis et al., 2005). Lewis et al. (2005) found the same result by expanding the space of unrooted tree topologies to include multifurcations. Our results show that this solution to the star-tree paradox extends to rooted trees.

### 3.1 Robustness of coalescent models that assume unlinked characters

Our finding that the multi-species coalescent model of Bryant et al. (2012) is robust to linked characters is consistent with previous simulations using species trees with one and two tips (Oaks, 2019; Oaks et al., 2019, 2020). Our simulation results show that this robustness extends to larger trees with multifurcations and shared divergences, and suggest that discarding data to avoid linked characters can have a worse effect on inference than violating the assumption of unlinked characters. This is consistent with the findings of Chifman and Kubatko (2014) that quartet inference of splits in multi-species coalescent trees from SNP data was also robust to the violation of the assumption that characters are unlinked.

### 3.2 Diversification of Philippine gekkonid lizards

How the 7,100 islands of the Philippines accumulated one of the highest concentrations of terrestrial biodiversity on Earth (Catibog-Sinha and Heaney, 2006; Brown and Diesmos, 2009; Heaney and Regalado, 1998; Brown et al., 2013) has been of interest to evolutionary biologists since the founding of biogeography (Wallace, 1869; Huxley, 1868; Dickerson, 1928; Diamond and Gilpin, 1983; Brown, 2016; Lomolino et al., 2016). Since the late Pliocene, the archipelago’s five major (and several minor) aggregate island complexes were repeatedly fragmented by interglacial rises in sea level into clusters of landmasses resembling today’s islands, followed by island fusion via land bridge exposure as sea levels fell during glacial periods (Haq et al., 1987; Rohling et al., 1998; Siddall et al., 2003; Miller et al., 2005; Spratt and Lisiecki, 2016). The repeated fragmentation-fusion cycles of this insular landscape has generated a prominent hypothesis to explain the high levels of terrestrial biodiversity across the Philippines (Inger, 1954; Heaney, 1985; Brown and Guttman, 2002; Evans et al., 2003; Heaney et al., 2005; Roberts, 2006; Linkem et al., 2010; Siler et al., 2010, 2011, 2012; Brown and Siler, 2014). However, there is growing evidence that (1) older tectonic processes (~30–5 mya) of precursor paleoislands (Jansa et al., 2006; Blackburn et al., 2010; Siler et al., 2012; Brown and Siler, 2014; Brown et al., 2016), (2) dispersal events from mainland source populations (Diamond and Gilpin, 1983; Brown and Guttman, 2002; Brown and Siler, 2014; Chan and Brown, 2017), (3) repeated colonizations among islands (Siler et al., 2011; Justiniano et al., 2015; Brown et al., 2016), and (4) fine-scale *in situ* isolating mechanisms (Heaney et al., 2011; Linkem et al., 2011; Siler et al., 2011, 2012; Hosner et al., 2013; Brown et al., 2015), have been important causes of diversification among and within many of the islands.

Oaks et al. (2019) found support for independent divergence times among inter-island pairs of *Cyrtodactylus* and *Gekko* populations from across the Philippines, suggesting that rare, over-water colonization, perhaps mediated by rafting on vegetation, might have been a more important mechanism of isolation than sea-level fragmentation in these gekkonid lizards. Our fully phylogenetic approach to this problem has allowed us to look for shared divergences across the full evolutionary history of extant populations in these clades, finding evidence for shared divergences that were missed by the pairwise approach. These results emphasize a pitfall of previous methods: choosing pairs of populations, for comparison under previous methods for inferring shared divergences (Hickerson et al., 2006; Huang et al., 2011; Oaks, 2014, 2019), was problematic in the sense that it was somewhat arbitrary and could miss more complex patterns of shared divergences in the shared ancestry of the taxa under study.

We recognize that our use of secondary calibrations to convert the timescale of the diversification of each genus into millions of years is error-prone, and should not be used to tie estimated shared divergences to specific geological or climatic events. However, given how recent most of the estimated shared divergences are for both *Cyrtodactylus* and *Gekko* (Figure 6; Table S3), it is unlikely the magnitude of error from our calibrations is great enough such that the true timing of these divergences would pre-date the late Pliocene. Thus, we conclude these estimated divergences are consistent with predictions of the model of diversification based on Plio-Pleistocene interglacial fragmentation of the islands. Given the numbers of shared or multifurcating divergence events estimated within this time frame are relatively low for both *Cyrtodactylus* and *Gekko* (6 and 5, respectively), and have weak to moderate support, our findings also are consistent with accumulating evidence that a number of complex processes of diversification have played important roles in shaping the distribution of life across the Philippines, not just paleo-island fragmentation.

A simultaneous analysis involving broader taxonomic sampling of Philippine gekkonids (*e.g., Gekko, Cyrtodactylus, Pseudogekko, Lepidodactylus*, and *Luperosaurus*; Wood, Jr. et al., 2020) would likely reveal support for an increased number of shared divergences across the archipelago, including older divergences predicted by geological processes that pre-date the Plio-Pleistocene interglacial fragmentations. When comparing our results between *Cyrtodactylus* and *Gekko*, we see some patterns suggestive of such shared diversification. For example, early divergences in both genera show patterns consistent with arrival into the archipelago, and subsequent diversification, via the Palawan Island Arc (Blackburn et al., 2010; Siler et al., 2012). Results from both genera support a pattern where a clade, sister to species endemic to the Palawan microcontinental block, began diversifying across the oceanic islands of the Philippines approximately 25–20 mya (Figure 6). Among the divergences estimated to have occurred within the last 2 my, there also appear to be regional consistencies in when and where lineages were diversifying in the Philippines, including population-level diversification for the widespread *Cyrtodactylus philippinicus* and *Gekko mindorensis* within and among the Mindoro and West Visayan faunal regions in the central Philippines (Figures 6, S14, & S15; Siler et al., 2012, 2014). Regardless of temporal concordance among divergences, the results of this work further support Philippine species within both focal clades having originated in the archipelago as a result of one or more faunal exchanges between oceanic portions of the Philippines associated historically with the Philippine mobile belt and the Palawan microcontinental block (Brown et al., 2013; Yumul et al., 2008).

Currently, broader taxonomic analyses are limited by a simplifying assumption of `phycoeval` that mutation rates are constant across the tree. We sought to minimize the effects of violations of this assumption by analyzing the two gekkonid genera separately. The Philippine species in each genus are closely related (the posterior mean root age in expected substitutions per site for *Cyrtodactylus* and *Gekko* was 0.012 and 0.013, respectively) and share similar natural histories, so an assumption of a similar rate of mutation across the populations we sampled within each genus seems reasonable. Future developments of `phycoeval` allowing the rate to vary across the phylogeny would be an obvious way to improve our current implementation and make it more generally applicable to a greater diversity of systems.

### 3.3 Future directions

Given that processes of co-diversification are of interest to fields as diverse as biogeography, epidemiology, and genome evolution, we hope the generalized tree model offers a statistical framework for studying these processes across the life sciences. To help achieve this, there are several ways to improve upon our current implementation of this approach. Allowing the generalized tree model and associated MCMC algorithms to be coupled with a diverse set of phylogenetic likelihood models is an obvious way to expand its applicability to more data types and systems. The independence of the tree model and MCMC algorithms from the likelihood function makes this relatively straightforward. Similarly, our approach can be extended to accommodate tips sampled through time (Stadler, 2010; Heath et al., 2014; Gavryushkina et al., 2016; Stadler et al., 2018) and “relaxed-clock” models (Drummond et al., 2006; Drummond and Suchard, 2010; Heath et al., 2011). The former would allow for fossil and epidemiological data, and the latter would allow it to be applied to diverse sets of taxa that are expected to vary in their rates of mutation.

As we alluded to above when discussing MCMC behavior, expanding the set of tree models to include all possible non-reticulating topologies with one to *N* – 1 divergence times could have important implications for the joint posterior distribution of phylogenetic models. We suggest posteriors that are rugged under a tree model with strictly independent and bifurcating divergences might be smoother under a generalized tree model, but more formal theoretical work to characterize this joint space is needed.

Lastly, the distribution we used over the generalized tree space (uniform over topologies with beta-distributed node heights) is motivated by mathematical convenience, rather than inspired by biological processes. Process-based models, like a generalized birth-death model, could provide additional insights. In addition to inferring phylogenies with shared or multifurcating divergences, process-based models would allow us to infer the macroevolutionary parameters that govern the rate of such divergences.

## 4 Methods

### 4.1 Generalized tree model

Let *T* represent a rooted, potentially multifurcating tree topology with *N* tips and *n*(*t*) internal nodes ***t*** = *t*_1_, *t*_2_, … *t_n_*(*t*), where *n*(*t*) can range from 1 (the “comb” tree) to *N* – 1 (fully bifurcating, independent divergences). Each internal node t is assigned to one divergence time *τ*, which it may share with other internal nodes in the tree. We will use ***τ*** = *τ*_1_, …, *τ_n_*_(*τ*)_ to represent *n*(*τ*) divergence times, where *n*(*τ*) can also range from 1 to *N* – 1, and every *τ* has at least one node assigned to it, and every node maps to a divergence time more recent than its parent (Figure S17).

To formalize a distribution across this space of generalized trees, we assume all possible topologies (*T*) are equally probable (see Figure S1 for an example of the sample space of topologies). We also assume the age of the root node follows a parametric distribution (e.g., a gamma distribution), and each of the other divergence times is beta-distributed between the present (*τ*_0_) and the height of the youngest parent of a node mapped to the divergence time (Figure S17). This was inspired by and related to the Dirichlet distribution on divergence times of Kishino et al. (2001), but we use beta distributions to make it easier to deal with the fact that under our generalized tree model, multiple nodes can be mapped to each divergence time. For additional flexibility, we allow a distribution to be placed on the alpha parameter of the beta distributions of all the non-root divergence times, which we denote as *α_τ_*.

### 4.2 Likelihood model

To perform Bayesian phylogenetic inference under the generalized tree model, it can be coupled with any function for calculating the probability of data evolving along a tree. This means it can be coupled with any data type and associated phylogenetic likelihood function. Even if the likelihood function does not explicitly accommodate multifurcations, these can be treated as a series of arbitrary bifurcations with branches of zero length to obtain the same likelihood of the tree.

Here, we couple the generalized tree model with a multi-species coalescent model that allows the likelihood of any species tree to be estimated directly from biallelic character data, while analytically integrating out all possible gene trees and character substitution histories along those gene trees. Below we give a brief overview of this model; for a full description of this likelihood model, please see Bryant et al. (2012), and see Oaks (2019) for a correction when only variable characters are analyzed.

#### 4.2.1 The data

From *N* species for which we wish to infer a phylogeny, we assume we have collected orthologous, biallelic genetic characters. By “biallelic”, we mean that each character has at most two states, which we refer to as “red” and “green” following Bryant et al. (2012). For each character from each species, we have collected *n* copies of the locus, *r* of which are copies of the red allele. We will use **n** and **r** to denote allele counts for one character from all *N* species; i.e., **n**, **r** = {(*n*_1_, *r*_1_), (*n*_2_, *r*_2_), … (*n_N_*, *r_N_*)}. We use **D** to represent these allele counts across all the characters.

#### 4.2.2 The evolution of characters

We assume each character evolved along a gene tree (*g*) according to a finite-characters, continuous-time Markov chain (CTMC) model, and the gene tree of each character is in-dependent of the others, conditional on the species tree (i.e., the characters are effectively unlinked). We use *u* and *v* to denote the relative rate of mutating from the red to green state and vice versa, respectively, as a character evolves along a gene tree, forward in time (Bryant et al., 2012; Oaks, 2019). Thus, *π* = *u*/(*u* + *v*) is the stationary frequency of the green state. We denote the overall rate of mutation as *μ*, which we assume is constant across the tree (i.e., a “strict clock”). Because evolutionary change is the product of *μ* and time, when *μ* = 1, time is measured in units of expected substitutions per character. If a mutation rate per character per unit of time is given, then time is measured in those units (e.g., generations or years).

#### 4.2.3 The evolution of gene trees

We assume the gene trees of each character branched according to a multi-species coalescent model within a single, shared, generalized species tree, where each branch *i* represents a population with a constant effective size 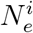 (Nielsen and Wakeley, 2001; Rannala and Yang, 2003; Liu and Pearl, 2007; Heled and Drummond, 2010; Bryant et al., 2012). We use ***N**_e_* to denote the effective population sizes for all branches in the generalized tree, with topology *T* and divergence times ***τ***; 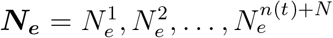 where *n*(*t*) + *N* is equal to the number of branches in the tree.

#### 4.2.4 The likelihood

Using the work of Bryant et al. (2012), we analytically integrate over all possible gene trees and character substitution histories to compute the likelihood of the species tree directly from all *m* biallelic characters under a multi-population coalescent model (Kingman, 1982a,b; Rannala and Yang, 2003),

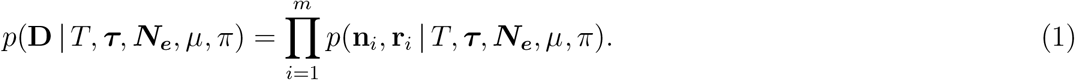

To accommodate multifurcations, we used recursion and Equation 19 of Bryant et al. (2012). This equation shows how to obtain the conditional probabilities at the bottom of an ancestral branch by merging the conditional probabilities at the top of its two descendant branches. At a multifurcation, we recursively apply Equation 19 of Bryant et al. (2012) to merge the conditional probabilities of each descendant branch in arbitrary order. We confirmed that this recursion returns an identical likelihood as treating the multifurcation as a series of bifurcations with zero-length branches.

### 4.3 Bayesian inference

The joint posterior probability distribution of the tree (with potential shared and multifurcating divergences) and other model parameters is

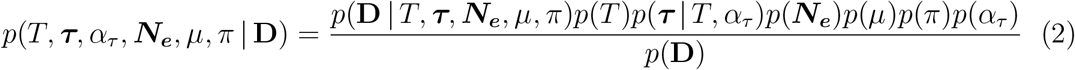

#### 4.3.1 Priors

We use the generalized tree distribution described above as the prior on the topology (*T*) and divergence times (***τ***). For all of our analyses below, we (1) set the alpha parameter of the beta distributions on non-root divergence times (*α_τ_*) to 1, (2) set the mutation rate (*μ*) to 1, so that time is in units of expected substitutions per character, (3) assume one gamma-distributed effective population size is shared across all the branches of the species tree, and (4) set the stationary frequencies of the two character states to be equal (π = 0.5), making our CTMC model of character evolution a two-state equivalent to the “JC69” model of nucleotide substitution (Jukes and Cantor, 1969).

### 4.4 Approximating the posterior of generalized trees

We use Markov chain Monte Carlo (MCMC) algorithms (Metropolis et al., 1953; Hastings, 1970; Green, 1995) to sample from the joint posterior in Equation 2. To sample across trees with different numbers of divergence times during the MCMC chain, we use reversible-jump MCMC (Green, 1995). We also use univariate and multivariate Metropolis-Hastings algorithms (Metropolis et al., 1953; Hastings, 1970) to update the divergence times and effective population sizes. See the Supporting Information for details and validations of our MCMC algorithms.

### 4.5 Software implementation

We implemented the models and algorithms above for approximating the joint posterior distribution of generalized trees, divergence times, and other model parameters in the software package `ecoevolity` Oaks (2019); Oaks et al. (2019, 2020). The `C++` source code for `ecoevolity` is freely available from https://github.com/phyletica/ecoevolity and includes an extensive test suite. From the `C++` source code, three command-line tools are compiled for generalized tree analyses: (1) `phycoeval`, for performing Bayesian inference under the model described above, (2) `simphycoeval` for simulating data under the model described above, and (3) `sumphycoeval` for summarzing the posterior samples of generalized trees collected by `phycoeval`. Documentation for how to install and use the software is available at http://phyletica.org/ecoevolity/. A detailed, version-controlled history of this project, including all of the data and scripts needed to produce our results, is available as a GitHub repository https://github.com/phyletica/phycoeval-experiments and was archived on zenodo (Oaks, 2021). We used multiple commits of `ecoevolity` for the analyses below, as we added features to the `sumphycoeval` tool (this history is documented in the project repository). However, all of our analyses can be replicated using Version 1.0.0 (Commit 2ed8d6ec) of `ecoevolity`.

### 4.6 Simulation-based analyses

#### 4.6.1 Methods used for all our simulations (unless noted)

We used `sumphycoeval` to simulate data sets of 50,000 biallelic characters from one diploid individual from nine species (i.e., two copies of each character sampled from each species). Except for our simulations of linked characters described below, the characters were unlinked (i.e., each character was simulated along an independent gene tree within the species tree). For all of our simulations and analyses, we constrained the branches of the species tree to share the same mutation-scaled, diploid effective population size (*N_e_ μ*), which we randomly drew from a gamma distribution with a shape of 20 and mean of 0.001. We used this distribution as the prior on *N_e_* in subsequent analyses of the simulated data sets. The mean of this distribution corresponds to an average number of differences per character between individuals of 0.004, which is comparable to estimates from genomic data from populations of zooplankton (Choquet et al., 2019), stickleback fish (Hohenlohe et al., 2010), humans (Auton et al., 2015), and the gecko species we analyze here (Oaks et al., 2019).

We analyzed each simulated data set under two models using `phycoeval`: the generalized tree model described above, which we denote as *M_G_*, and an otherwise equivalent model that is constrained to the space of trees with independent, bifurcating divergences (i.e., trees with *N* – 1 divergence times), which we denote as *M_IB_*. For both *M_G_* and *M_IB_*, we used a gamma-distributed prior on the age of the root node with a shape of 10 and mean of 0.2. For each data set we ran four independent MCMC chains for 15,000 generations, sampling every 10 generations, and retaining the last 1000 samples of each chain to approximate the posterior (4000 total samples). For each generation, nine (equal to the number of tips) MCMC moves are randomly selected in proportion to specified weights, some of which automatically call other moves after finishing to improve mixing. Each chain started from a random bifurcating topology with no shared divergences, and the root age and other divergence times drawn randomly from their respective prior distributions.

From the 4000 posterior samples collected for each simulated dataset, we used `sumphycoeval` to calculate the mean and 95% credible intervals of the root age, tree length, effective population size, and the number of divergence times, and to summarize the frequency of sampled topologies, splits, nodes, and shared divergences. We define a split as a branch in the tree that “splits” the tips of the tree into two non-overlapping subsets; those that do and do not descend from the branch. We define a node as a split with a particular set of splits that descend from it; this is necessary to summarize the frequency of multifurcations. We also used `sumphycoeval` to calculate the distance between every sampled tree and the true tree using the square root of the sum of squared differences in branch lengths (Robinson and Foulds, 1979; Kuhner and Felsenstein, 1994). To assess convergence and mixing of the chains, we used `sumphycoeval` to calculate the average standard deviation of split frequencies (ASDSF; Lakner et al., 2008) across the four chains with a minimum split frequency threshold of 10%, as well as the potential scale reduction factor (PSRF; the square root of Equation 1.1 in Brooks and Gelman, 1998) and effective sample size (ESS; Gong and Flegal, 2016) of the log likelihood, root age, tree length, and effective population size.

#### 4.6.2 Simulations on fixed trees

We used `simphycoeval` to simulate 100 data sets on two fixed trees with 9 species, one with shared and multifurcating divergences (Figure 2A) and the other with only bifurcating, independent divergences (Figure 3A). We analyzed each simulated data set under models *M_G_* and *M_IB_*, both with and without constant characters; for the latter we specified for `phycoeval` to correct the likelihood for only sampling variable characters (Bryant et al., 2012; Oaks, 2019).

To explore the improved MCMC mixing under the *M_G_* model for data sets simulated on trees with shared or multifurcating divergences (see results), we re-ran the analyses under the *M_IB_* on the 100 data sets simulated on the tree shown in Figure 2A. In these re-analyses under *M_IB_*, we ran the MCMC chain twice as long (30,000 generations versus 15,0000) while sampling half as frequently (every 20 generations versus 10), and started each chain with the correct tree.

#### 4.6.3 Simulations on random trees

Using `simphycoeval`, we also simulated 100 data sets on trees randomly drawn from the prior distributions of the *M_G_* and *M_IB_* models. As above, we analyzed each simulated data set with and without constant characters under the *M_G_* and *M_IB_* models. We used MCMC to sample trees randomly from the prior distributions of both models. More specifically, we used `simphycoeval` to (1) randomly assemble a strictly bifurcating tree with no shared divergences times, (2) run an MCMC chain of topology changing moves for a specified number of generations (we used 1000), and (3) draw the root age, other divergence times, and the effective population sizes randomly from their respective prior distributions. For each MCMC generation, nine (equal to the number of tips) topology changing moves were randomly selected in proportion to specified weights.

Due to the nested beta (uniform) distributions on non-root divergence times, some trees sampled from *M_G_* and *M_IB_* will have all or most of the divergence times close to zero. This happens when one of the oldest non-root divergences is randomly assigned a time near zero. For example, the trees shown in Figure S16A–C all have eight independent, bifurcating divergences. Given such trees, it is nearly impossible to differentiate independent divergences with a finite data set. It is also not clear what an investigator would want `phycoeval` to infer given a true tree like Figure S16A; whereas eight independent divergences is technically correct in the synthetic world of nested beta distributions, it seems an unlikely biological explanation. To avoid such extreme scenarios, we rejected any trees that had divergences times closer than 0.001 substitutions per site. This resulted in 61 and 201 trees being rejected in order to obtain 100 trees under the *M_G_* and *M_IB_* models, respectively. Despite this arbitrary filtering threshold, challenging tree shapes remained in our sample for simulations. For example, see the trees in Figure S16D–F, all with eight independent, bifurcating divergences.

#### 4.6.4 Simulations of linked characters

The likelihood model above assumes characters are unlinked (i.e., they evolved along gene trees that are independent of one another conditional on the species tree). To assess the effect on inference of violating this assumption, we repeated the simulations and analyses above (for both fixed and random trees), but simulated 500 loci of 100 linked characters each (i.e. for each locus, 100 characters evolved along a shared gene tree). We used `simphycoeval` to simulate these data sets in two ways: (1) all 50,000 characters are simulated and retained, and (2) only (at most) one variable character is retained for each locus. For the latter data sets, characters are unlinked, but only (at most) 500 characters, all variable, are sampled. We analyzed all of these data sets under both the *M_G_* and *M_IB_* models. For data sets with only variable characters, we corrected the likelihood for not sampling constant characters (Bryant et al., 2012; Oaks, 2019).

### 4.7 Inference of shared divergences in Philippine gekkonids

We applied our new approach to two genera of geckos, *Gekko* and *Cyrtodactylus*, sampled across the Philippine Islands. We used the RADseq data of Oaks et al. (2019) available on the NCBI Sequence Read Archive (Bioproject PRJNA486413, SRA Study SRP158258).

#### 4.7.1 Assembling alignments

We used ipyrad (Version 0.9.43; Eaton and Overcast, 2020) to assemble the RADseq reads into loci for both genera. All of the scripts and ipyrad parameter files we used to assemble the data are available in our gekkonid project repository (https://github.com/phyletica/gekgo) archived on Zenodo (Oaks and Wood, Jr., 2021), and the ipyrad settings are listed in Table S5. Using `pycoevolity` (Version 0.2.9; Commit 217dbeea; Oaks, 2019), we converted the ipyrad alignments into nexus format, and in the process, removed sites that had more than two character states. The final alignment for *Cyrtodactylus* contained 1702 loci and 155,887 characters from 27 individuals, after 567 characters with more than two states were removed. The final alignment for *Gekko* contained 1033 loci and 94,612 characters from 26 individuals, after 201 characters with more than two states were removed. Both alignments had less than 1% missing characters. The assembled data matrices for *Cyrtodactylus* and *Gekko* are available in our project repository (https://github.com/phyletica/phycoeval-experiments) and the data associated with specimens are provided in Tables S1 & S2.

#### 4.7.2 Phylogenetic analyses

When analyzing the *Cyrtodactylus* and *Gekko* character matrices with `phycoeval`, we (1) fixed stationary state frequencies to be equal (π = 0.5), (2) set the mutation rate (*μ*) to 1 so that divergence times are in units of expected substitutions per site, (3) used an exponentially distributed prior with a mean of 0.01 for the age of the root, (4) set *α_τ_* = 1 so that non-root divergence times are uniformly distributed between zero and the age of the youngest parent node, and (5) assumed a single diploid effective population size (*N_e_*) shared across the branches of the tree with a gamma-distributed prior. For the gamma prior on *N_e_*, we used a shape of 2.0 and mean of 0.0005 for *Cyrtodactylus*, and a shape of 4.0 and mean of 0.0002 for *Gekko*, based on estimates of Oaks et al. (2019) from the same and related species.

For both genera, we ran 25 independent MCMC chains for 15,000 generations, sampling the state of the chain every 10 generations. In each generation, `phycoeval` attempts *N* MCMC moves (27 and 26 for *Cyrtodactylus* and *Gekko*, respectively) randomly selected in proportion to specified weights, some of which automatically call other moves after finishing to improve mixing. For 20 of the chains, we specified for `phycoeval` to start from the “comb” topology (*n*(*τ*) = 1). For the remaining five chains, we had `phycoeval` start with a random bifurcating topology with no shared divergences (*n*(*τ*) = *N* – 1).

We used `sumphycoeval` to summarize the sampled values of all parameters and the frequency of sampled topologies, splits, nodes, and shared divergences. To assess convergence and mixing, we used `sumphycoeval` to calculate the average standard deviation of split frequencies (ASDSF; Lakner et al., 2008) and the potential scale reduction factor (PSRF; the square root of Equation 1.1 in Brooks and Gelman, 1998) and effective sample size (ESS; Gong and Flegal, 2016) of all parameters across all 25 MCMC chains. We present these convergence statistics in Table S4.

To plot the trees, we used `sumphycoeval` to scale the branch lengths of all the sampled trees so that the posterior mean root age was 23.07 million years for *Cyrtodactylus* (Grismer et al., 2022) and 33.76 million years for *Gekko*; the latter age is based on a time-calibrated phylogenetic estimate from another data set that is being prepared for publication. Our goal in scaling the branch lengths to millions of years is not to test whether shared divergence events correspond with the onset of specific interglacial periods, but rather to see if shared divergences fall within the general time frame predicted by a model of sea-level-driven diversification (i.e., within the last ≈4 million years).

## Data availability

We recorded a detailed history of all our analyses for this project in a version-controlled repository, which is publicly available at github.com/phyletica/phycoeval-experiments and archived on Zenodo (doi.org/10.5281/zenodo.5162056; Oaks, 2021). The `C++` source code for `ecoevolity` is freely available from github.com/phyletica/ecoevolity, with detailed documentation and tutorials available at phyletica.org/ecoevolity. All of the scripts and parameter files we used to assemble the gecko datasets are available in our gekkonid project repository (github.com/phyletica/gekgo) that is archived on Zenodo (doi.org/10.5281/zenodo.5162085; Oaks and Wood, Jr., 2021). The gecko sequence reads are available on the NCBI Sequence Read Archive (Bioproject PRJNA486413, SRA Study SRP158258).

## 5 Acknowledgments

We thank Mark Holder for helpful advice with Hastings ratios and modeling the distribution on divergence times. This work was supported by funding provided to JRO from the National Science Foundation (NSF grant number DEB 1656004). The computational work was made possible by the Auburn University (AU) Hopper and Easley Clusters supported by the AU Office of Information Technology and a grant of high-performance computing resources and technical support from the Alabama Supercomputer Authority. Our gecko sampling was amassed with NSF support for fieldwork (EF-0334952, DEB 073199 and 0743491 to RMB; and 0804115 to CDS) and Fulbright grants to CDS. This paper is contribution number 949 of the Auburn University Museum of Natural History.

## Supporting Information

### 1 Figures referenced in main text

**Figure S1.**
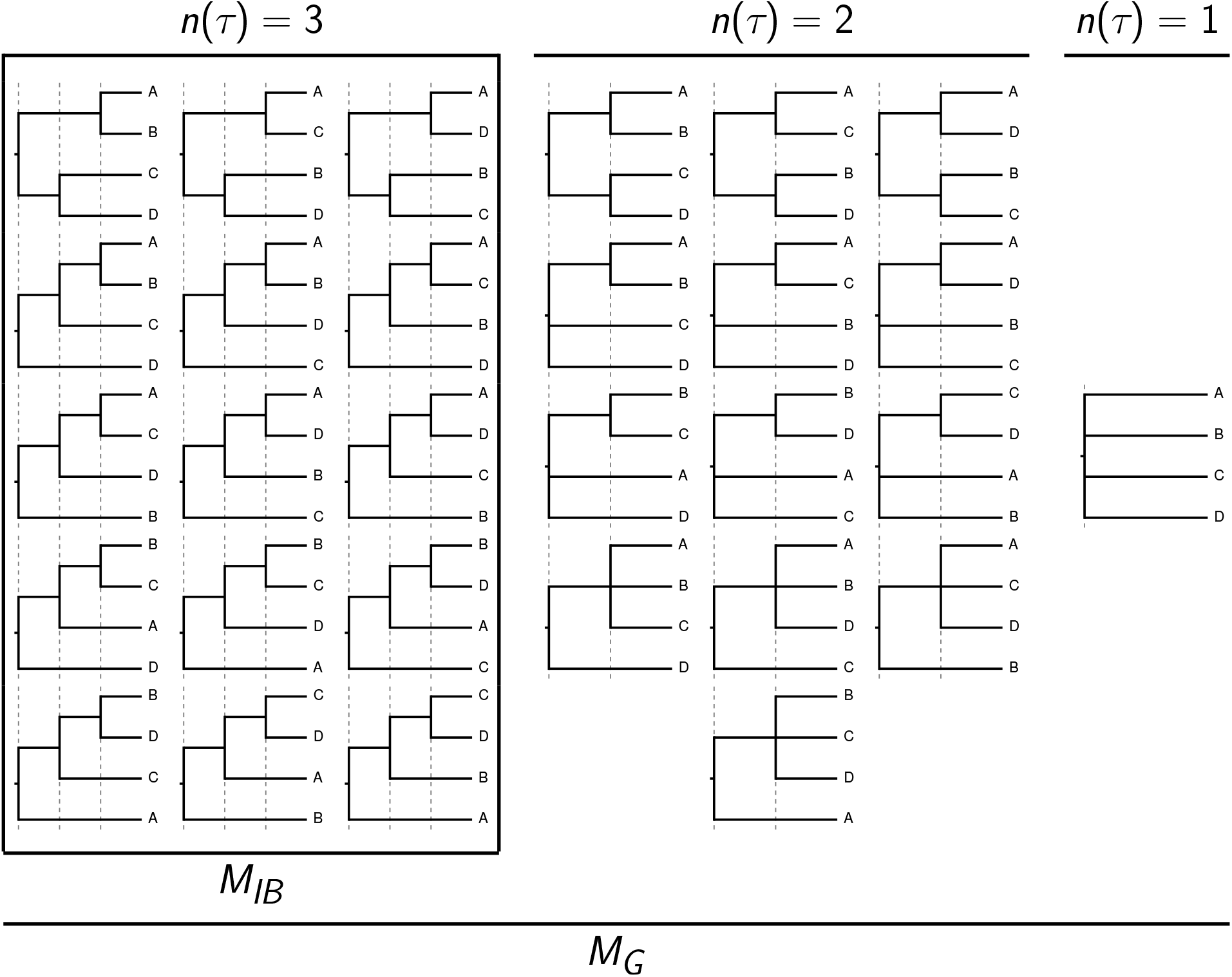
An example of the expanded sample space of topologies for the generalized tree distribution (*M_G_*) versus a tree model that assumes independent, bifurcating divergences (*M_IB_*). With four tips, there are 15 topologies for a model, all with three divergence times (within box). The *M_G_* model has 14 additional topologies with fewer than three divergence times. Notice the internal nodes are not labeled; i.e., the rank, or order, of non-nested internal nodes do not matter. *E.g*., in the topology at the top-left, regardless of whether A & B or C & D diverge first, it is the same topology. However, if A & B and C & D diverge at the same time, this is a different topology (or tree model) with one fewer divergence-time parameter (the topology in the upper-left of the *n*(*τ*) = 2 section). Trees plotted using Gram (Version 4.0.0; Foster, 2018) and the P4 phylogenetic toolkit (Version 1.4 5742542; Foster, 2004).

**Figure S2.**
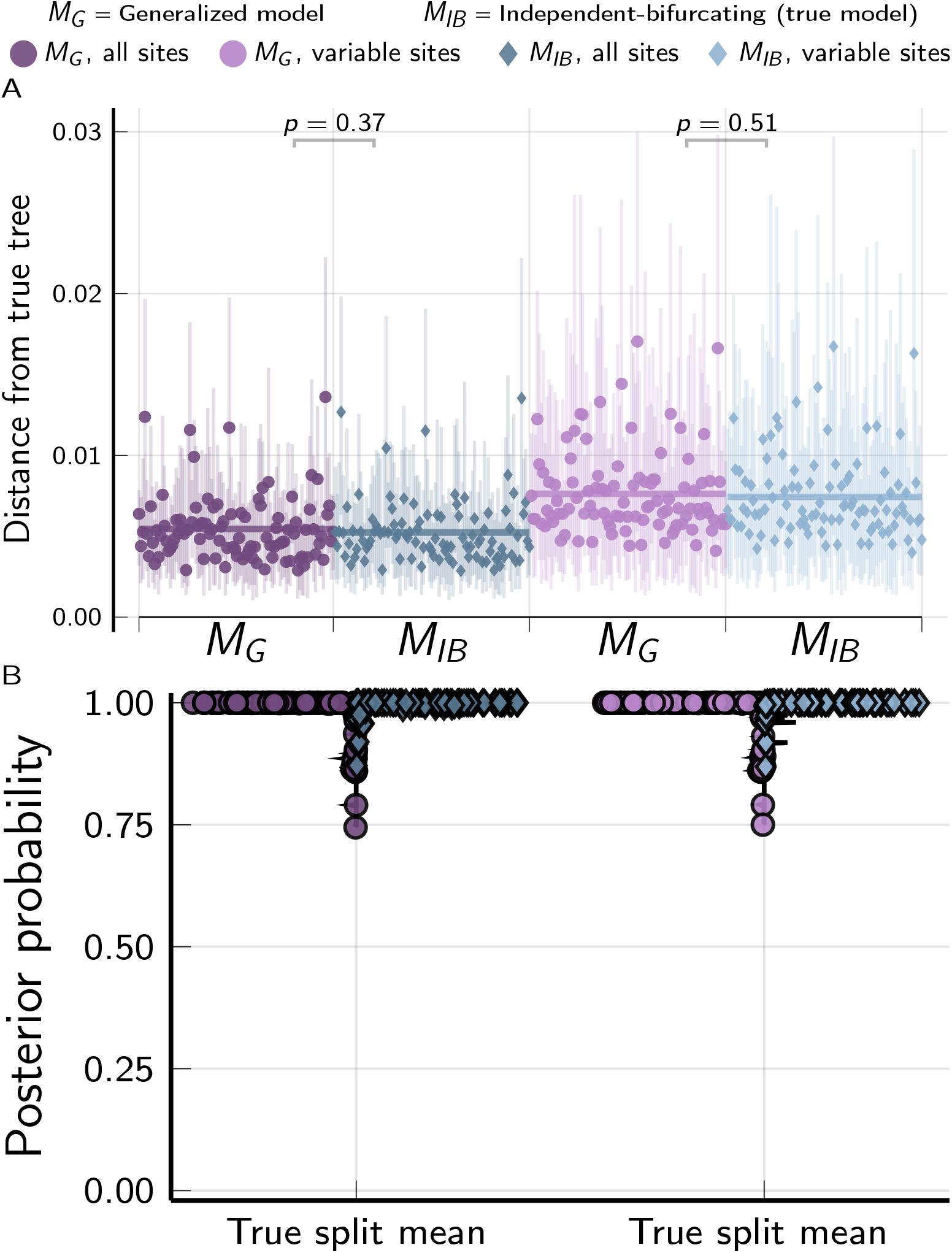
The performance of the *M_G_* and *M_IB_* tree models when applied to 100 data sets, each with 50,000 biallelic characters simulated on species trees randomly drawn from the *M_IB_* tree distribution. (A) The square root of the sum of squared differences in branch lengths between the true tree and each posterior tree sample (Kuhner and Felsenstein, 1994); the point and bars represent the posterior mean and equal-tailed 95% credible interval, respectively. P-values are shown for Wilcoxon signed-rank tests (Wilcoxon, 1945) comparing the paired differences in tree distances between methods. (B) Violin plots comparing the mean posterior probabilities of true splits for each of the 100 simulated trees. For each simulation, the mutation-scaled effective population size (*N_e_μ*) was drawn from a gamma distribution (shape = 20, mean = 0.001) and shared across all the branches of the tree; this distribution was used as the prior in analyses. Plots created using the PGFPlotsX (Version 1.2.10, Commit 1adde3d0; Carlsson and Papp, 2021) backend of the Plots (Version 1.5.7, Commit f80ce6a2; Breloff, 2021) package in Julia (Version 1.5.4; Bezanson et al., 2017).

**Figure S3.**
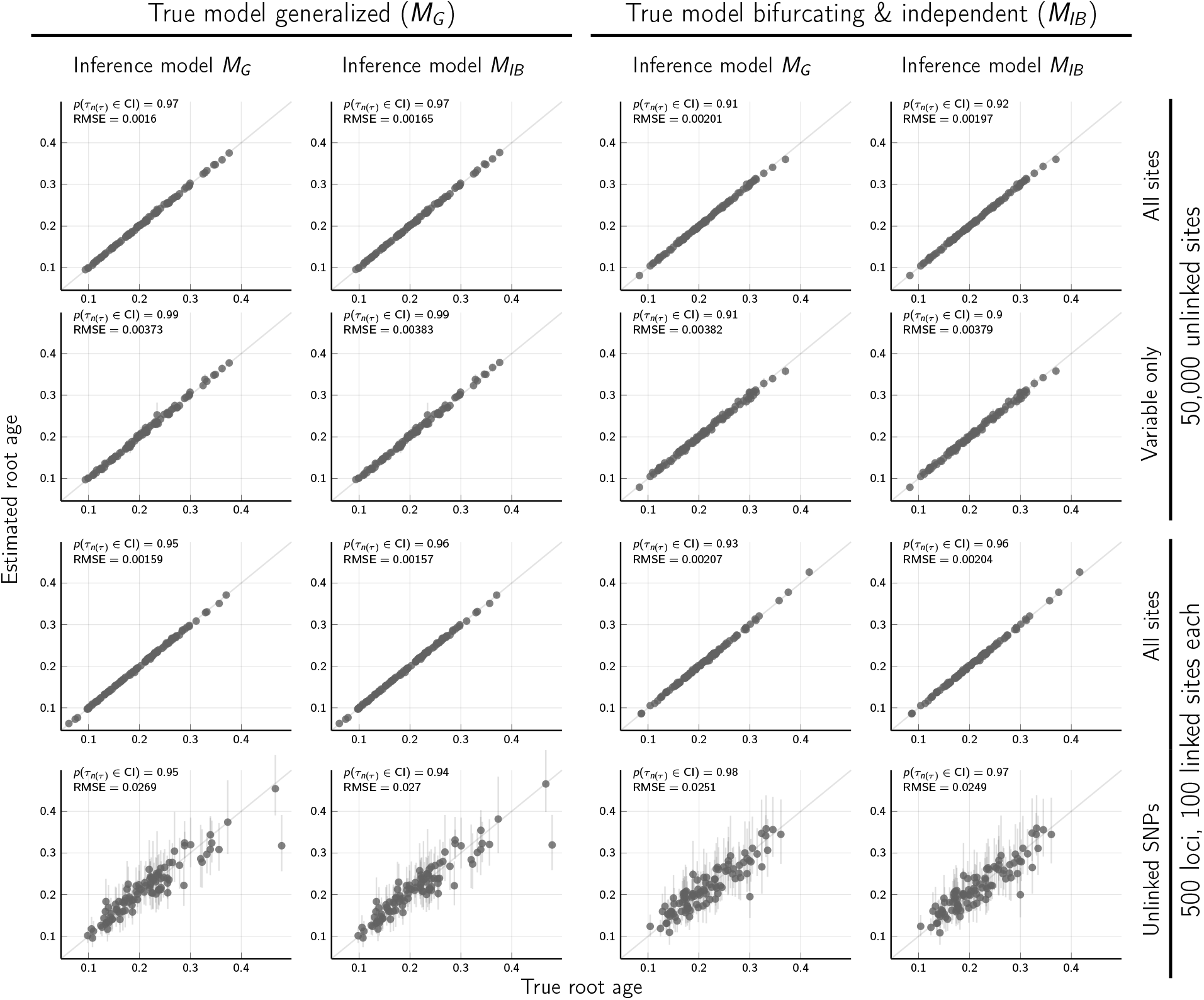
The accuracy and precision of the *M_G_* and *M_IB_* models at estimating the age of the root (in expected subsitutions per site) from data sets with 50,000 biallelic characters simulated on species trees randomly drawn from the *M_G_* and *M_IB_* tree distributions. Each plotted circle and associated error bars represent the posterior mean and 95% credible interval. Estimates for which the potential-scale reduction factor was greater than 1.2 (Brooks and Gelman, 1998) or the effective sample size was less than 200 are highlighted in red. Plots created using the PGFPlotsX (Version 1.2.10, Commit 1adde3d0; Carlsson and Papp, 2021) backend of the Plots (Version 1.5.7, Commit f80ce6a2; Breloff, 2021) package in Julia (Version 1.5.4; Bezanson et al., 2017).

**Figure S4.**
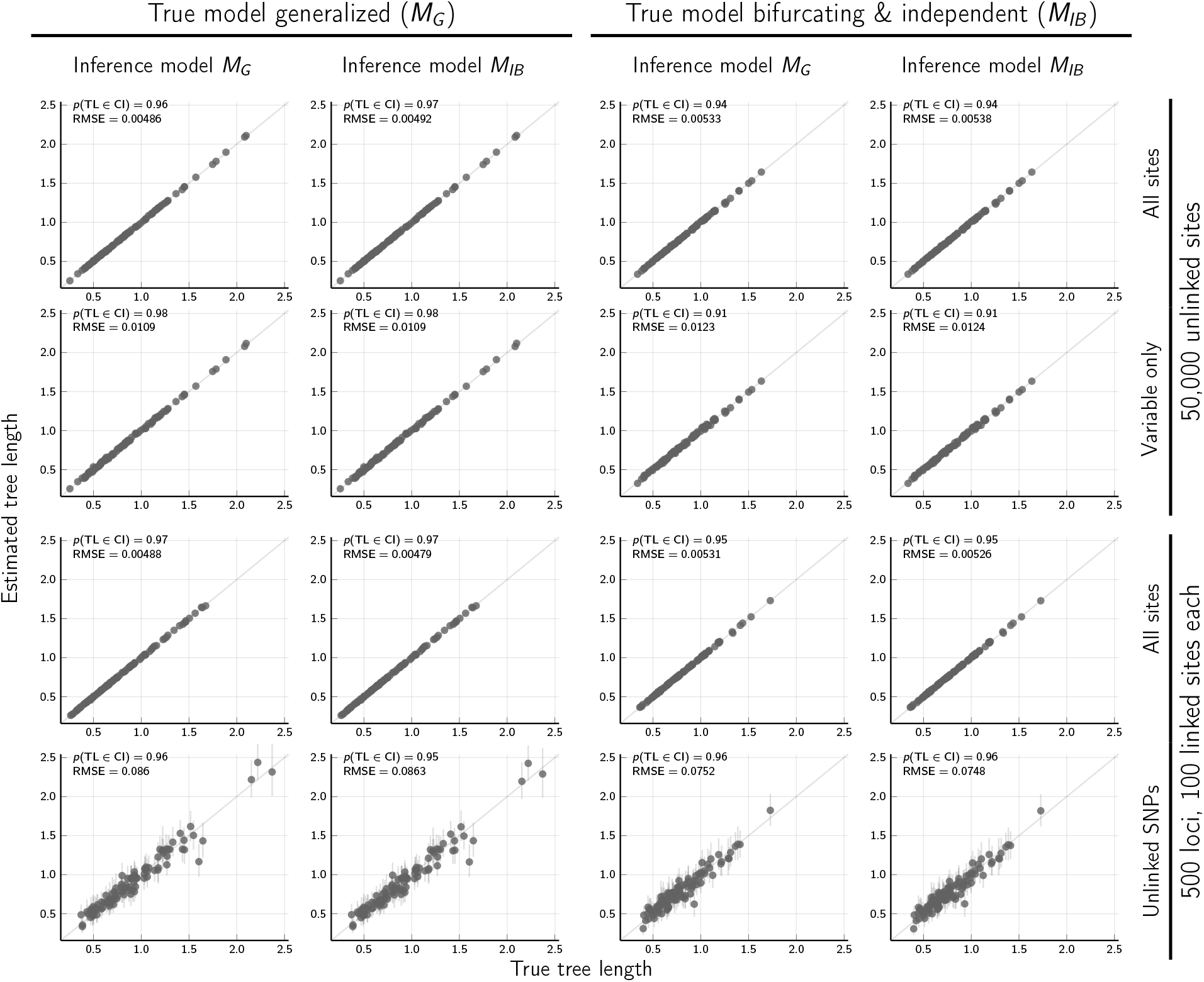
The accuracy and precision of the *M_G_* and *M_IB_* models at estimating the tree length (the sum of all branch lengths in units of expected substitutions per site) from data sets with 50,000 biallelic characters simulated on species trees randomly drawn from the *M_G_* and *M_IB_* tree distributions. Each plotted circle and associated error bars represent the posterior mean and 95% credible interval. Estimates for which the potential-scale reduction factor was greater than 1.2 (Brooks and Gelman, 1998) or the effective sample size was less than 200 are highlighted in red. Plots created using the PGFPlotsX (Version 1.2.10, Commit 1adde3d0; Carlsson and Papp, 2021) backend of the Plots (Version 1.5.7, Commit f80ce6a2; Breloff, 2021) package in Julia (Version 1.5.4; Bezanson et al., 2017).

**Figure S5.**
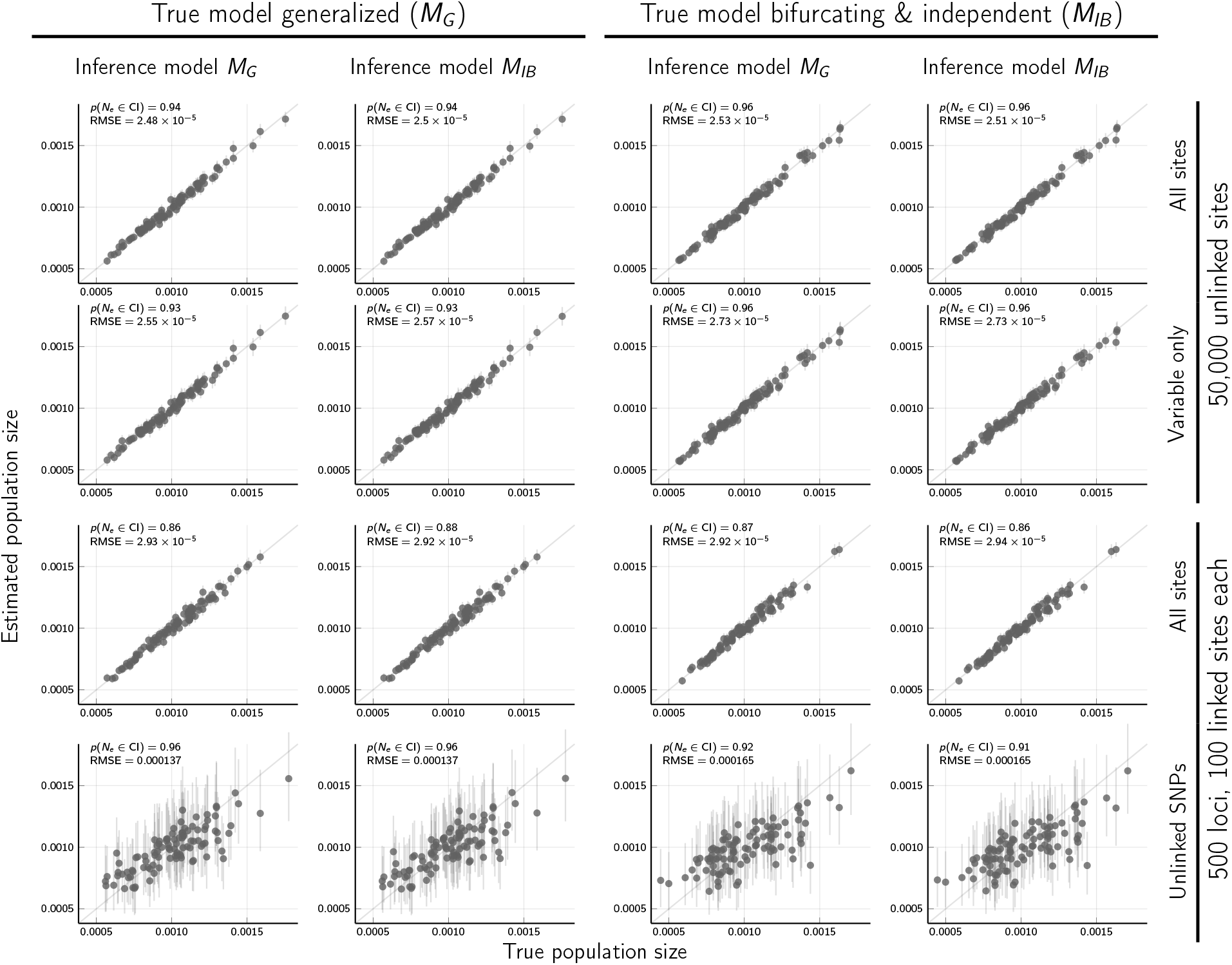
The accuracy and precision of the *M_G_* and *M_IB_* models at estimating the effective population size (*N_e_μ*) across the tree from data sets with 50,000 biallelic characters simulated on species trees randomly drawn from the *M_G_* and *M_IB_* tree distributions. Each plotted circle and associated error bars represent the posterior mean and 95% credible interval. Estimates for which the potential-scale reduction factor was greater than 1.2 (Brooks and Gelman, 1998) or the effective sample size was less than 200 are highlighted in red. Plots created using the PGFPlotsX (Version 1.2.10, Commit 1adde3d0; Carlsson and Papp, 2021) backend of the Plots (Version 1.5.7, Commit f80ce6a2; Breloff, 2021) package in Julia (Version 1.5.4; Bezanson et al., 2017).

**Figure S6.**
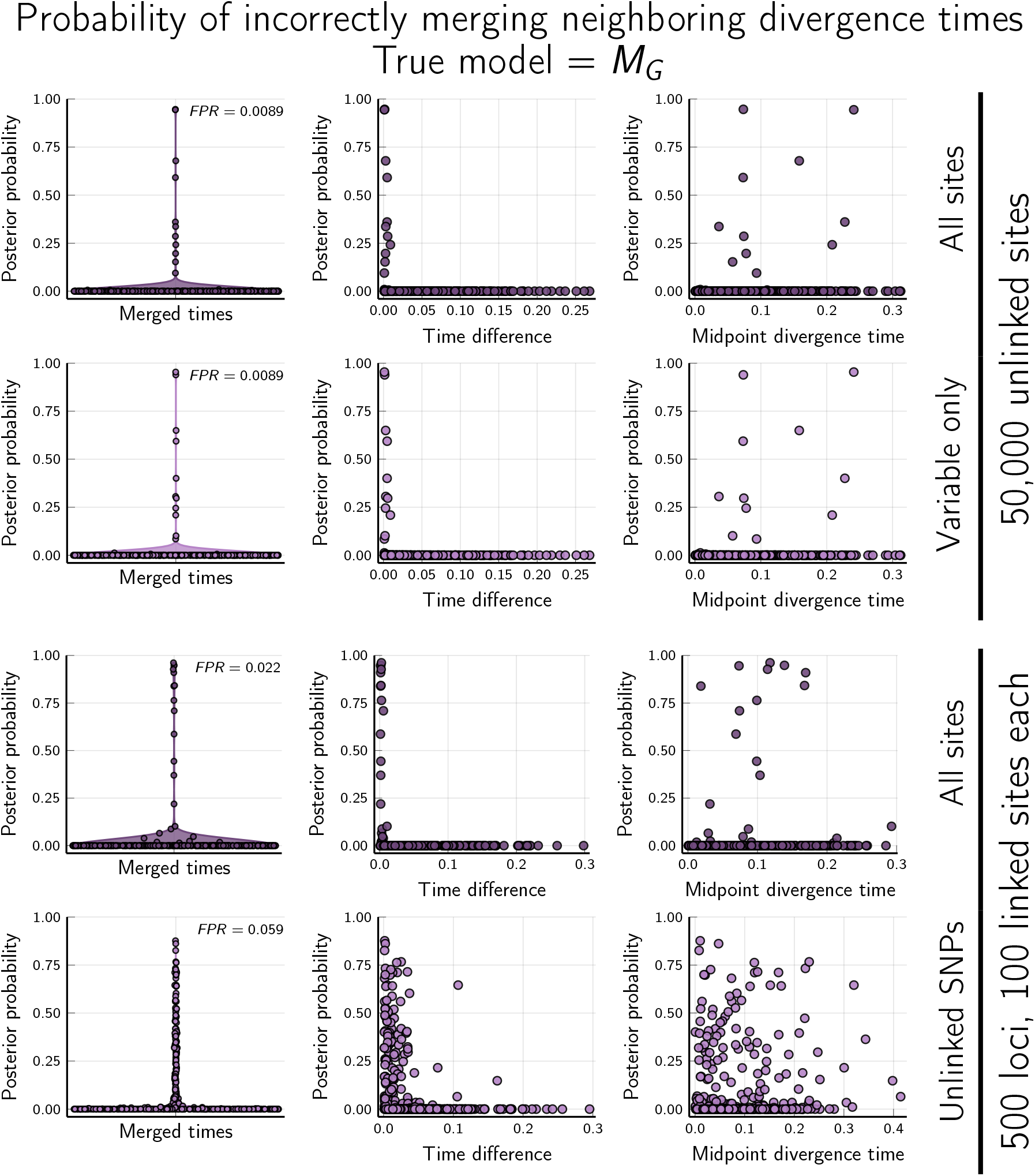
The *M_G_* model has a low false positive rate (FPR; the proportion of incorrectly merged divergence times with a posterior probability > 0.5) when applied to data simulated on trees drawn from *M_G_* with all (Row 1) or only variable (Row 2) unlinked characters, and all characters from linked loci (Row 3). Support for incorrectly merged divergence times is high only when the difference between the times is small (Column 2), and is not correlated with the age of the merged nodes (right). (Column 3; *P* = 0.109, 0.106, 0,053, and 0.068, from top to bottom for a t-test that Pearson’s correlation coefficient = 0 using all points with posterior probability > 0). When data sets with linked loci are reduced to only one variable site per locus (Row 4), the FPR increases (left) and precision decreases (right). The top row is the same as Figure 5A–C. Time units are expected substitutions per site. Plots created using the PGFPlotsX (Version 1.2.10, Commit 1adde3d0; Carlsson and Papp, 2021) backend of the Plots (Version 1.5.7, Commit f80ce6a2; Breloff, 2021) package in Julia (Version 1.5.4; Bezanson et al., 2017).

**Figure S7.**
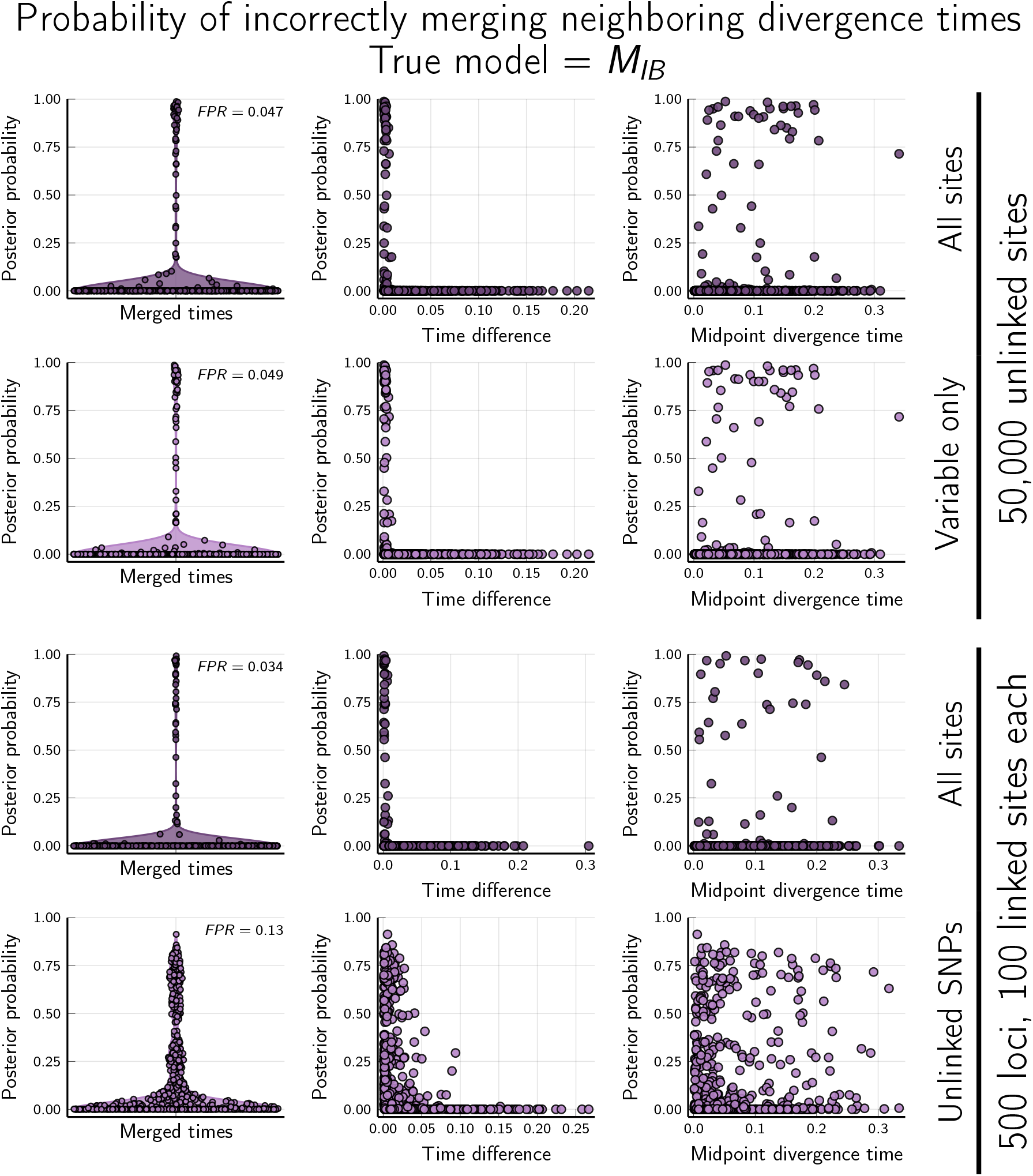
The *M_G_* model has a low false positive rate (FPR; the proportion of incorrectly merged divergence times with a posterior probability > 0.5) when applied to data simulated on trees drawn from *M_IB_* (no shared or multifurcating divergences) with all (Row 1) or only variable (Row 2) unlinked characters, and all characters from linked loci (Row 3). Support for incorrectly merged divergence times is high only when the difference between the times is small (Column 2), and is not correlated with the age of the merged nodes (right). (Column 3; *P* = 0.25, 0.29, 0,11, and 0.11, from top to bottom for a t-test that Pearson’s correlation coefficient = 0 using all points with posterior probability > 0). When data sets with linked loci are reduced to only one variable site per locus (Row 4), the FPR increases (left) and precision decreases (right). The top row is the same as Figure 5D–F. Time units are expected substitutions per site. Plots created using the PGFPlotsX (Version 1.2.10, Commit 1adde3d0; Carlsson and Papp, 2021) backend of the Plots (Version 1.5.7, Commit f80ce6a2; Breloff, 2021) package in Julia (Version 1.5.4; Bezanson et al., 2017).

**Figure S8.**
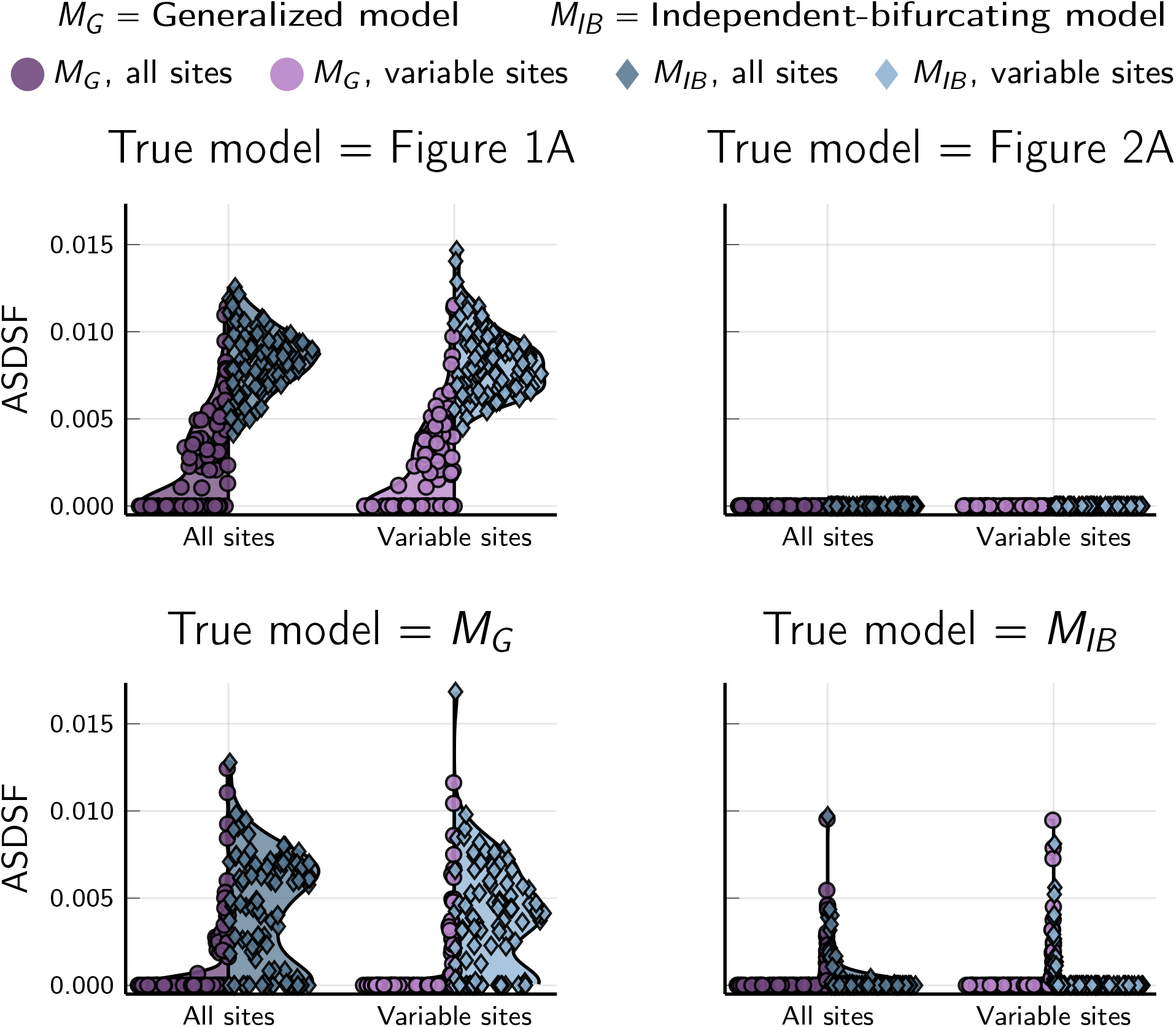
Markov chain Monte Carlo (MCMC) sampling yielded better convergence and mixing among chains under the generalized tree model when there were shared or multifurcating divergences. Convergence and mixing of sampled trees is summarized using the average standard deviation of split frequencies (ASDSF; Lakner et al., 2008) across the four chains of each analysis; smaller standard deviations across chains indicate better sampling behavior. Plots created using the PGFPlotsX (Version 1.2.10, Commit 1adde3d0; Carlsson and Papp, 2021) backend of the Plots (Version 1.5.7, Commit f80ce6a2; Breloff, 2021) package in Julia (Version 1.5.4; Bezanson et al., 2017).

**Figure S9.**
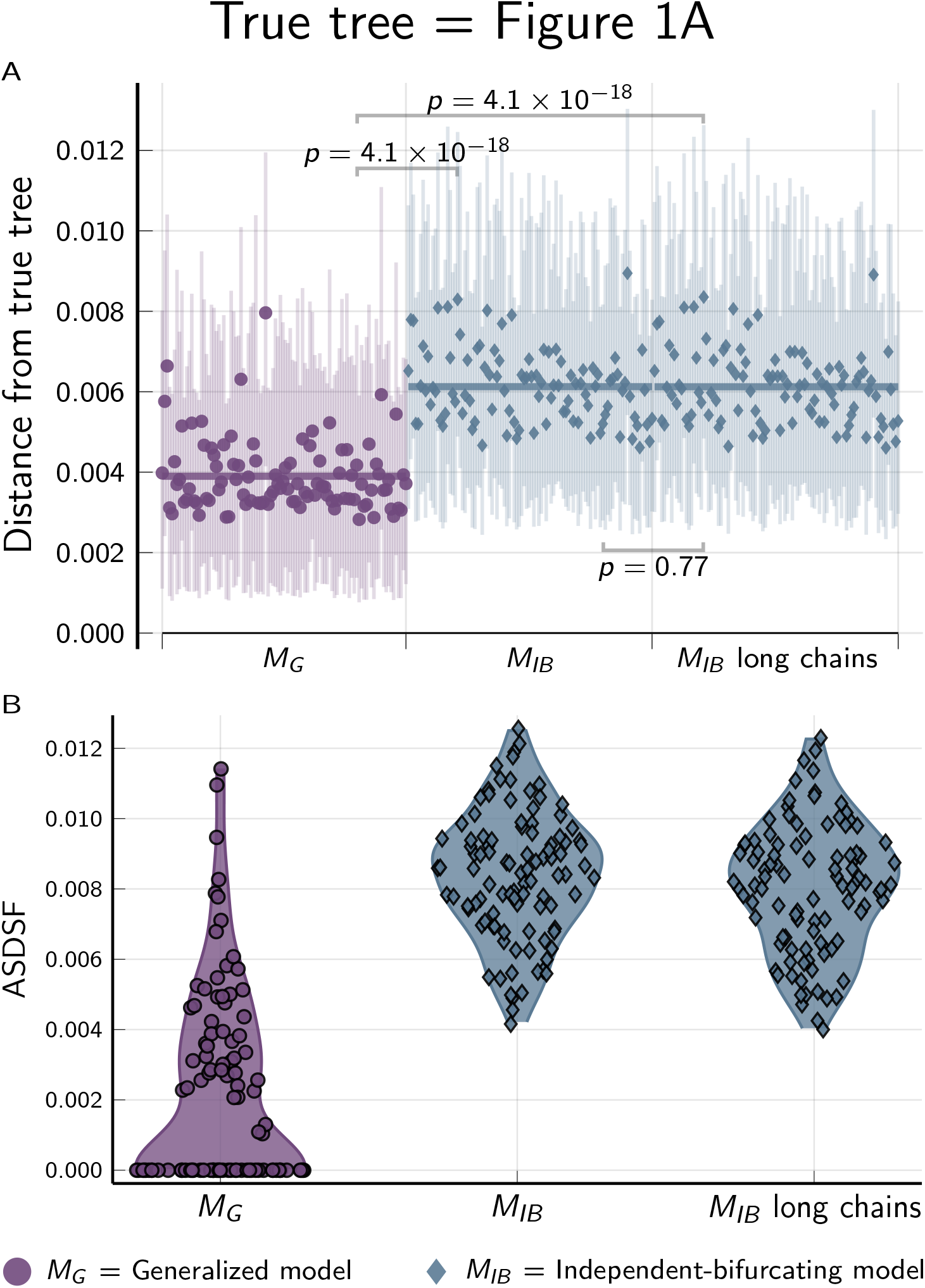
The (A) accuracy and (B) Markov chain Monte Carlo (MCMC) sampling efficiency of the *M_G_* tree model is better than *M_IB_*, even when the MCMC chains for *M_IB_* are started with the correct tree and are run twice as long and sampled half as frequently (“long chains”; 30,000 generations, sampling every 20). Results from 100 data sets simulated on the species tree shown in Figure 2A. (A) The square root of the sum of squared differences in branch lengths between the true tree and each posterior tree sample (Kuhner and Felsenstein, 1994); the point and bars represent the posterior mean and equal-tailed 95% credible interval, respectively. P-values are shown for Wilcoxon signed-rank tests (Wilcoxon, 1945) comparing the paired differences in tree distances between methods. (B) Violin plots of the average standard deviation of split frequencies (ASDSF; Lakner et al., 2008,; smaller is better) across the four chains of each analysis. The *M_G_* and *M_IB_* results (left and center) are the same shown in Figure 2B and Figure S8. Plots created using the PGFPlotsX (Version 1.2.10, Commit 1adde3d0; Carlsson and Papp, 2021) backend of the Plots (Version 1.5.7, Commit f80ce6a2; Breloff, 2021) package in Julia (Version 1.5.4; Bezanson et al., 2017).

**Figure S10.**
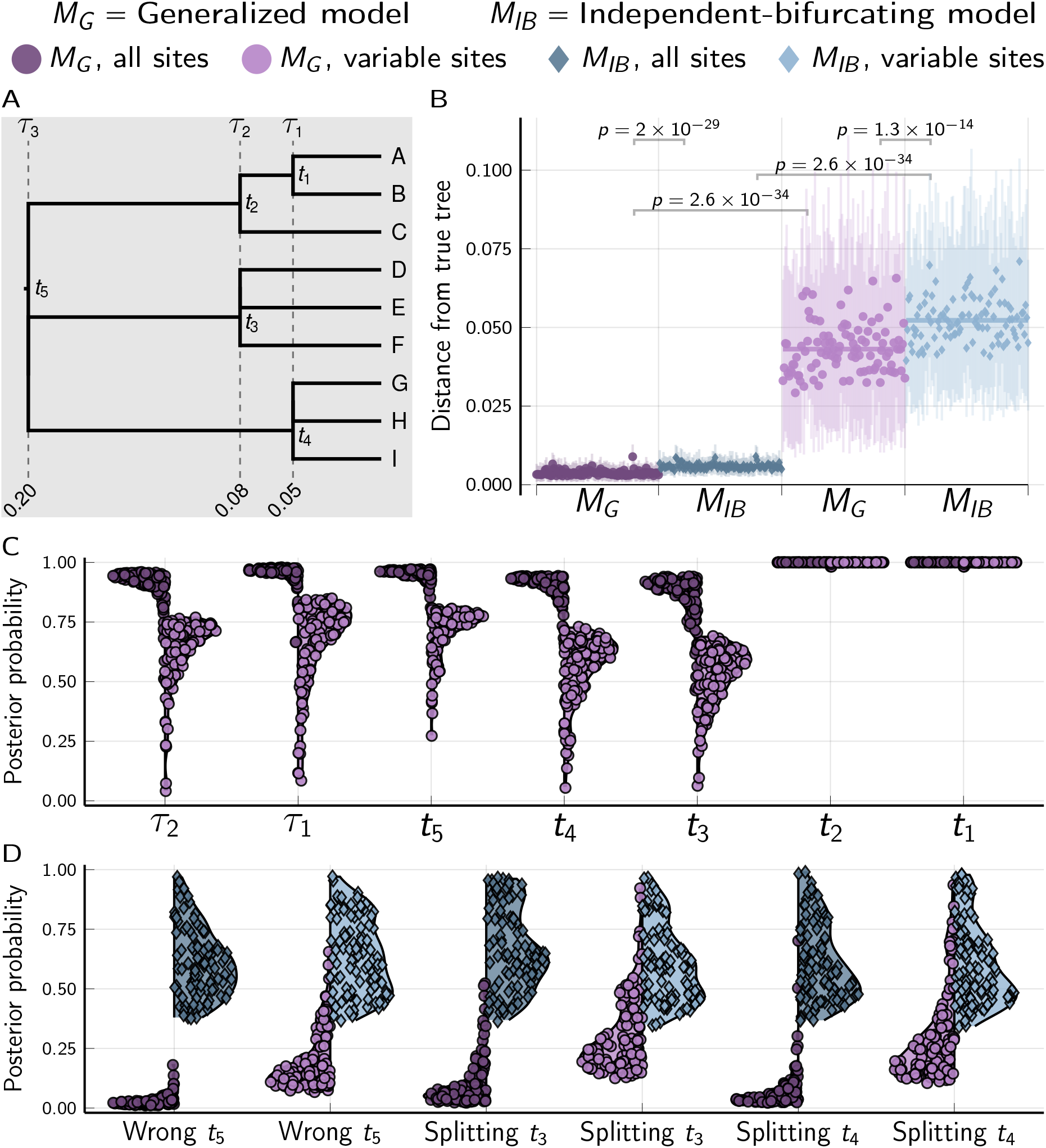
Results of analyses of 100 data sets with linked loci simulated along the species tree shown in (A). Each simulated data set comprised 500 loci with 100 biallelic characters. P-values are shown for Mann-Whitney U tests Mann and Whitney (1947) comparing the differences in tree distances between methods. For each simulation, the mutation-scaled effective population size (*N_e_μ*) was drawn from a gamma distribution (shape = 20, mean = 0.001) and shared across all the branches of the tree; this distribution was used as the prior in analyses. Tree plotted using Gram (Version 4.0.0, Commit 02286362; Foster, 2018) and the P4 phylogenetic toolkit (Version 1.4, Commit d9c8d1b1; Foster, 2004). Other plots created using the PGFPlotsX (Version 1.2.10, Commit 1adde3d0; Carlsson and Papp, 2021) backend of the Plots (Version 1.5.7, Commit f80ce6a2; Breloff, 2021) package in Julia (Version 1.5.4; Bezanson et al., 2017).

**Figure S11.**
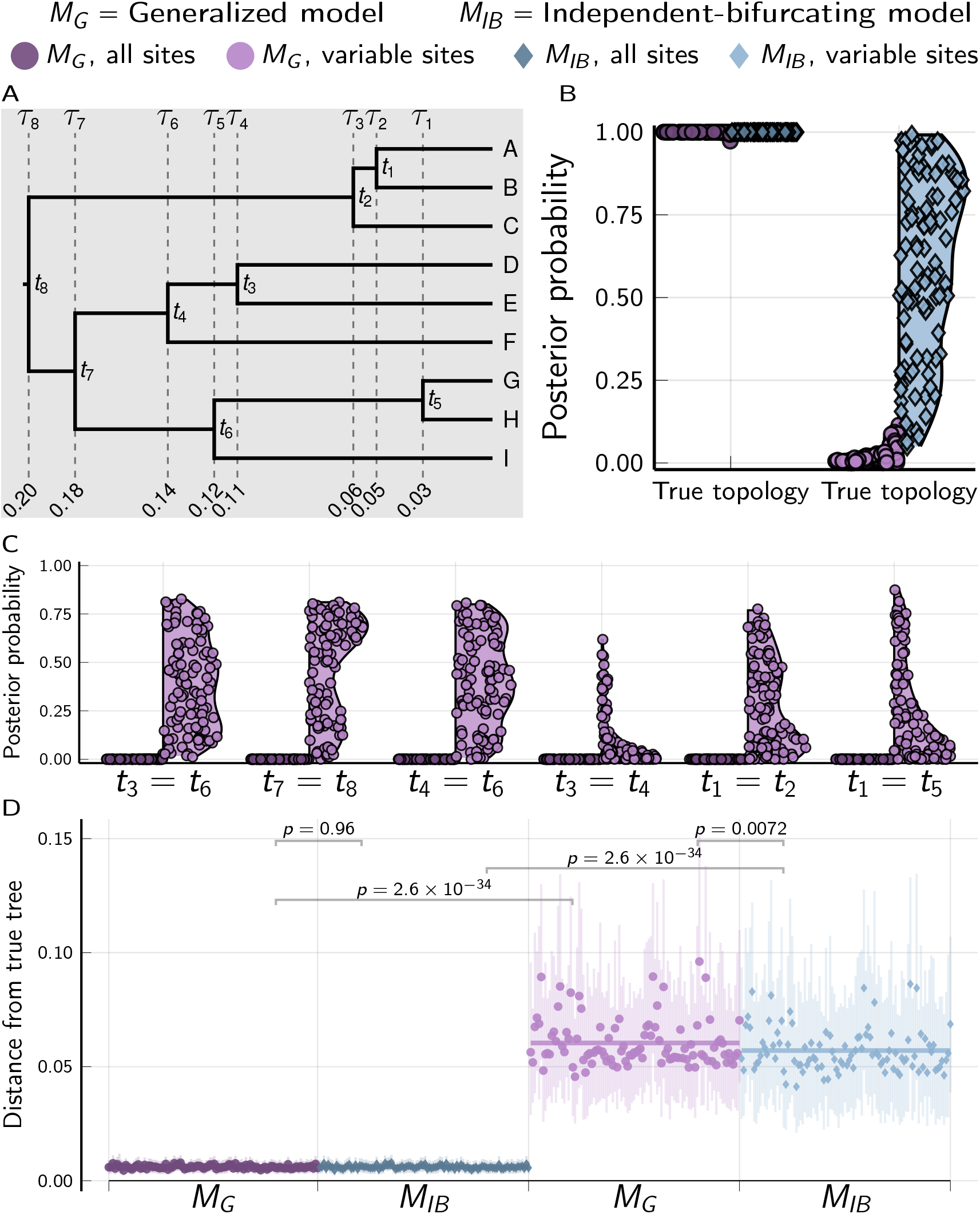
Results of analyses of 100 data sets with linked loci simulated along the species tree shown in (A). Each simulated data set comprised 500 loci with 100 biallelic characters. P-values are shown for Mann-Whitney U tests (Mann and Whitney, 1947) comparing the differences in tree distances between methods. For each simulation, the mutation-scaled effective population size (*N_e_μ*) was drawn from a gamma distribution (shape = 20, mean = 0.001) and shared across all the branches of the tree; this distribution was used as the prior in analyses. Tree plotted using Gram (Version 4.0.0, Commit 02286362; Foster, 2018) and the P4 phylogenetic toolkit (Version 1.4, Commit d9c8d1b1; Foster, 2004). Other plots created using the PGFPlotsX (Version 1.2.10, Commit 1adde3d0; Carlsson and Papp, 2021) backend of the Plots (Version 1.5.7, Commit f80ce6a2; Breloff, 2021) package in Julia (Version 1.5.4; Bezanson et al., 2017).

**Figure S12.**
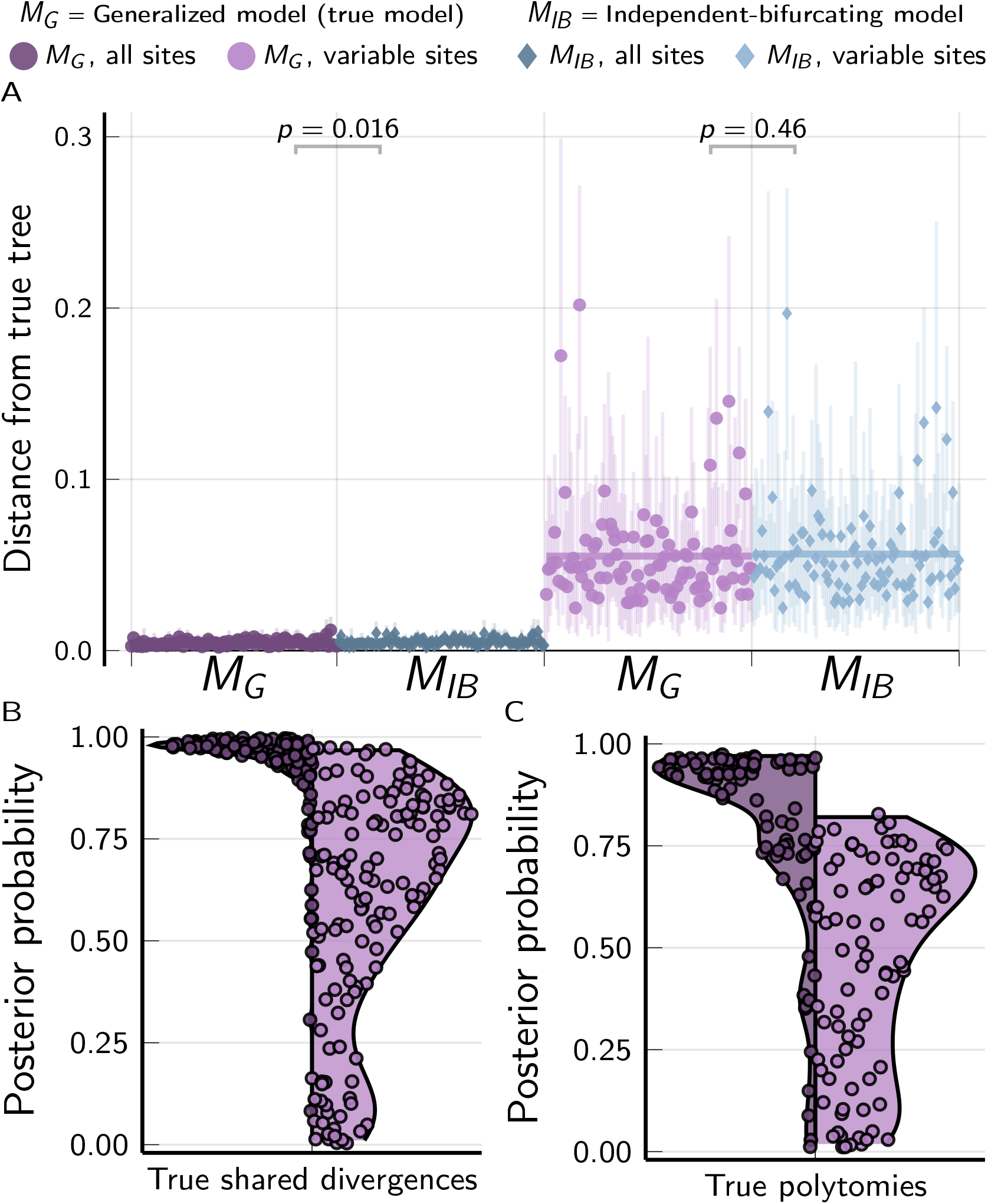
The performance of the *M_G_* and *M_IB_* tree models when applied to 100 data sets with 500 loci (each with 100 linked characters) simulated on species trees randomly drawn from the *M_G_* tree distribution. For each simulation, the mutation-scaled effective population size (*N_e_μ*) was drawn from a gamma distribution (shape = 20, mean = 0.001) and shared across all the branches of the tree; this distribution was used as the prior in analyses. Plots created using the PGFPlotsX (Version 1.2.10, Commit 1adde3d0; Carlsson and Papp, 2021) backend of the Plots (Version 1.5.7, Commit f80ce6a2; Breloff, 2021) package in Julia (Version 1.5.4; Bezanson et al., 2017).

**Figure S13.**
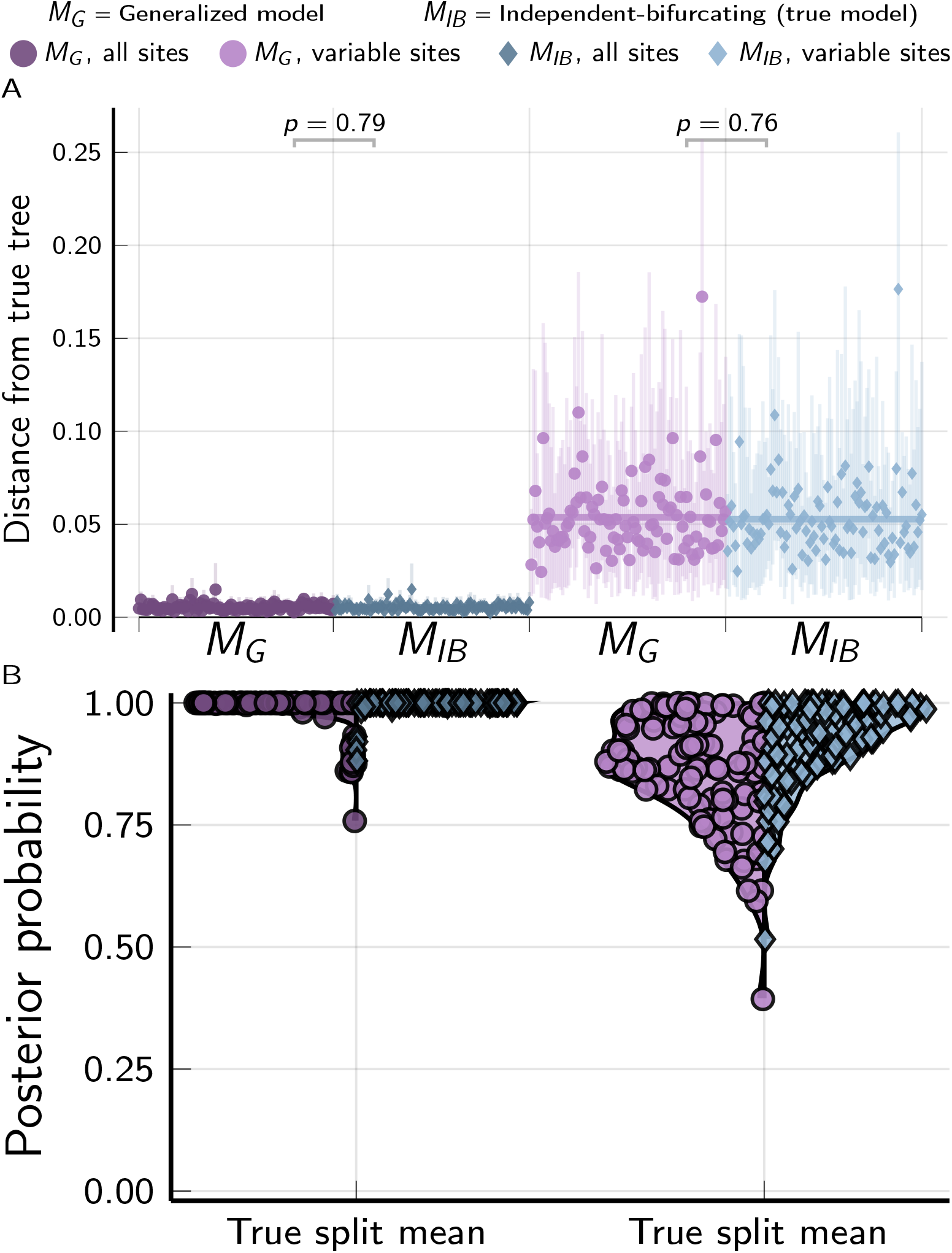
The performance of the *M_G_* and *M_IB_* tree models when applied to 100 data sets with 500 loci (each with 100 linked characters) simulated on species trees randomly drawn from the *M_IB_* tree distribution. For each simulation, the mutation-scaled effective population size (*N_e_μ*) was drawn from a gamma distribution (shape = 20, mean = 0.001) and shared across all the branches of the tree; this distribution was used as the prior in analyses. Plots created using the PGFPlotsX (Version 1.2.10, Commit 1adde3d0; Carlsson and Papp, 2021) backend of the Plots (Version 1.5.7, Commit f80ce6a2; Breloff, 2021) package in Julia (Version 1.5.4; Bezanson et al., 2017).

**Figure S14.**
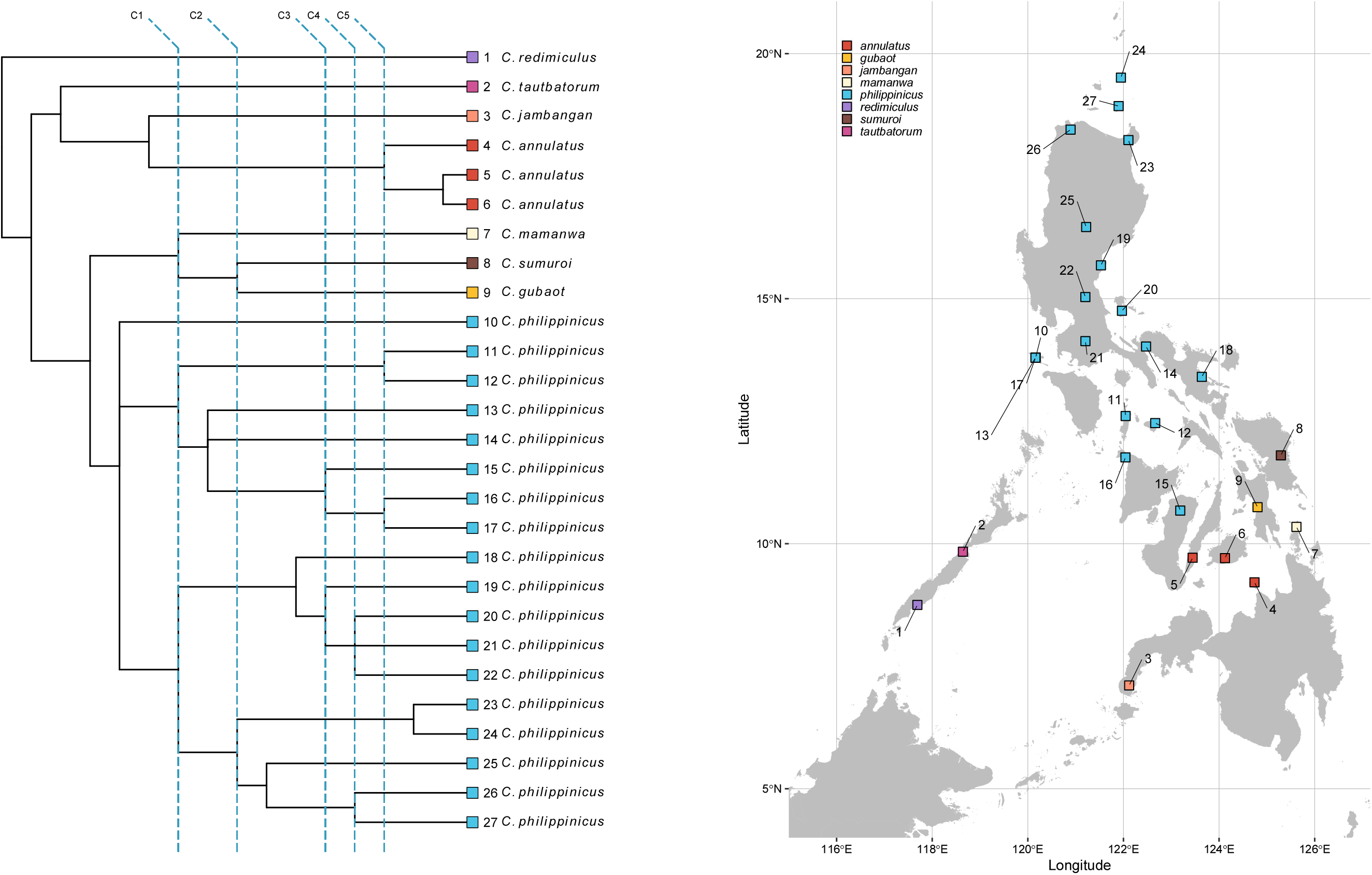
The maximum *a posteriori* (MAP) topology from Figure 6A for *Cyrtodactylus* shown without branch lengths to make it clearer which clades are involved. Shared divergences indicated by dashed lines, with labels shown along the top that correspond to rows in Table S3, where the divergences are summarized. Created using ggplot2 (v3.3.5; Wickham, 2016), ggtree (v3.1.0; Yu et al., 2017), treeio (v1.17.0; Wang et al., 2019), cowplot (v1.1.1; Wilke, 2020), and ggrepel (v0.9.1; Slowikowski, 2020). Link to nexus-formatted annotated MAP tree: *[Cyrtodactylus](https://raw.githubusercontent.com/phyletica/phycoeval-experiments/master/gekkonid-output/map-tree-cyrt-nopoly.nex)*.

**Figure S15.**
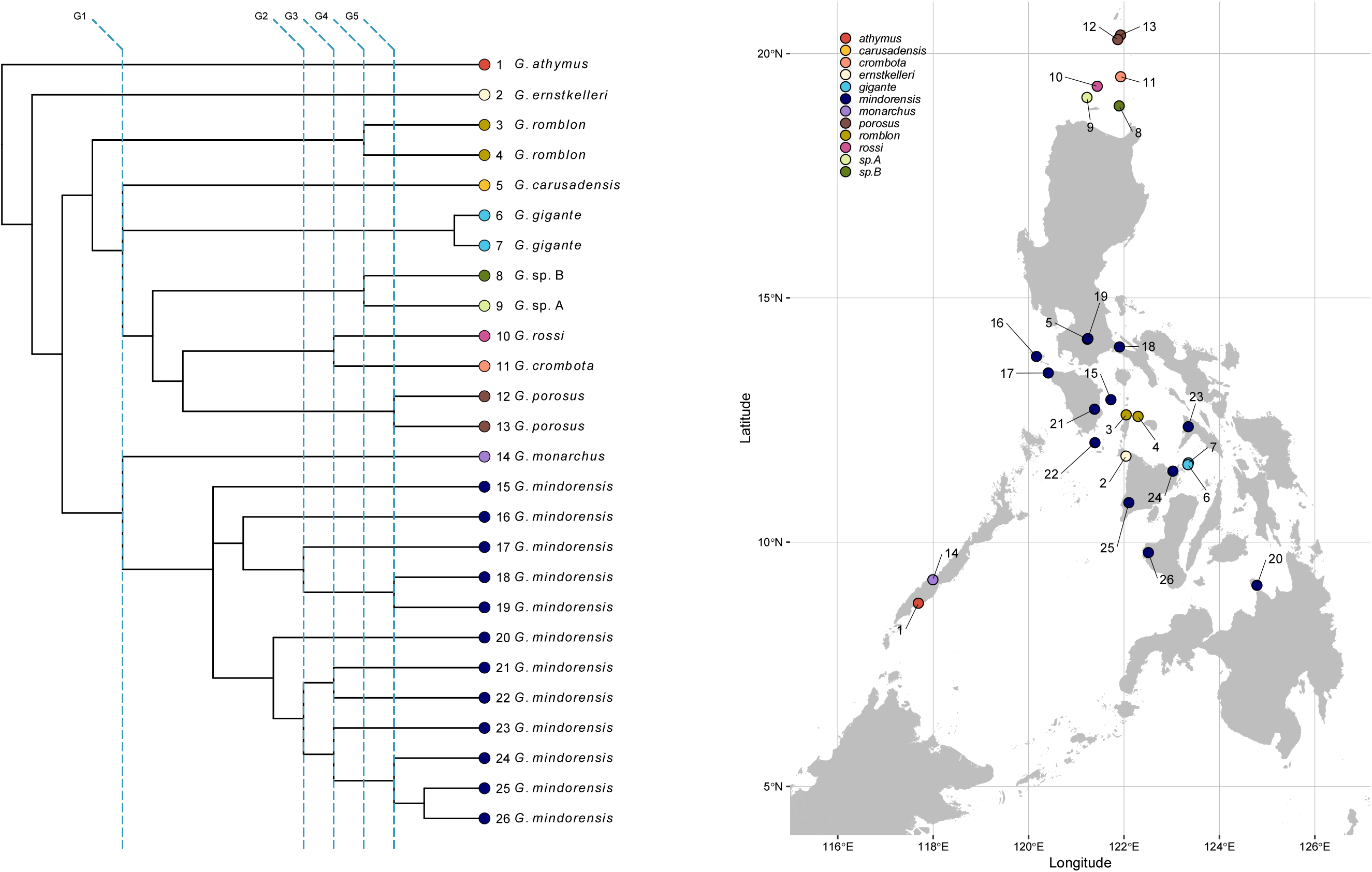
The maximum *a posteriori* (MAP) topology from Figure 6C for *Gekko* shown without branch lengths to make it clearer which clades are involved. Shared divergences indicated by dashed lines, with labels shown along the top that correspond to rows in Table S3, where the divergences are summarized. Created using ggplot2 (v3.3.5; Wickham, 2016), ggtree (v3.1.0; Yu et al., 2017), treeio (v1.17.0; Wang et al., 2019), cowplot (v1.1.1; Wilke, 2020), and ggrepel (v0.9.1; Slowikowski, 2020). Link to nexus-formatted annotated MAP tree: *[Gekko](https://raw.githubusercontent.com/phyletica/phycoeval-experiments/master/gekkonid-output/map-tree-gekko-nopoly.nex)*.

**Figure S16.**
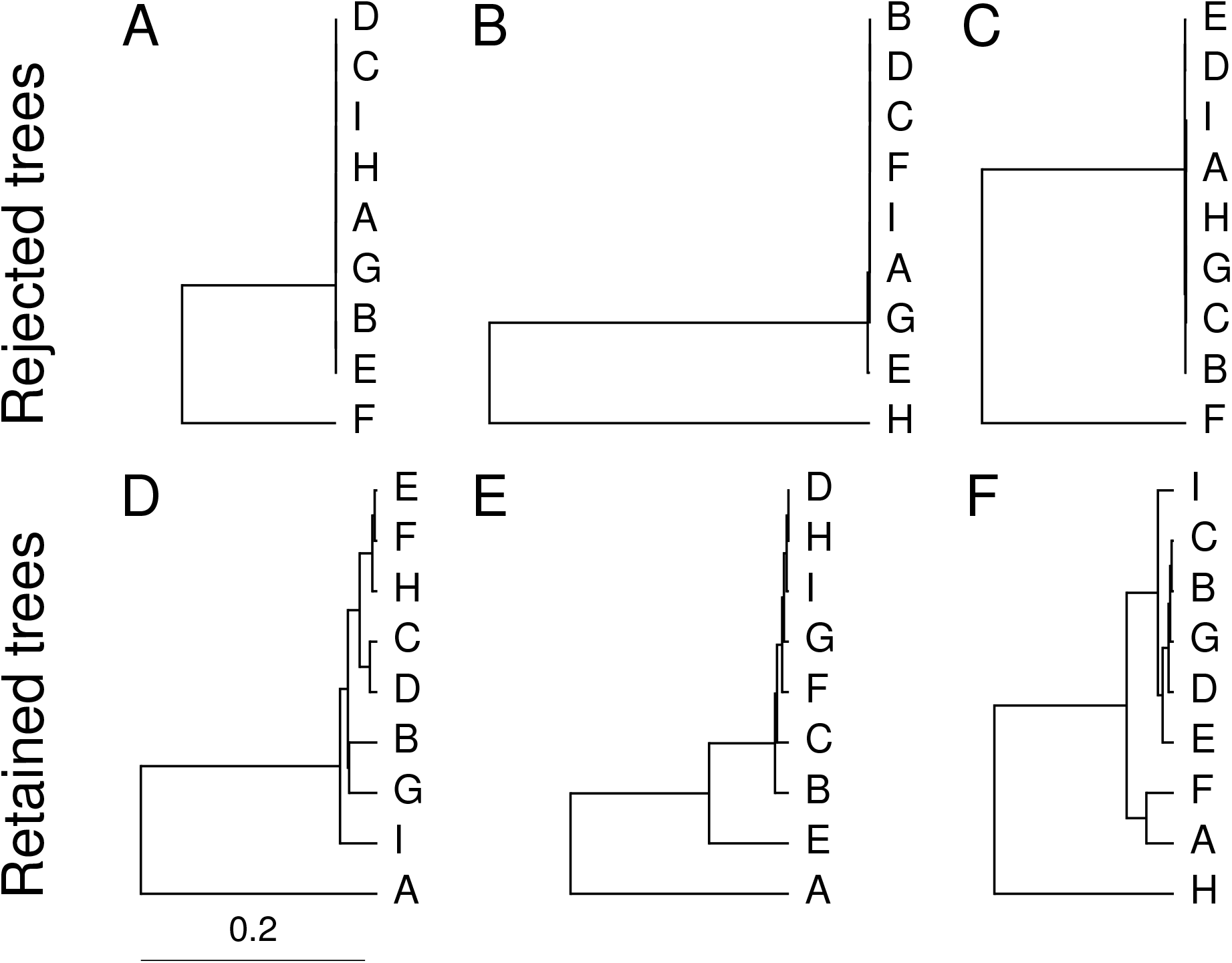
Examples of trees (all with 8 independent, bifurcating divergences) rejected (top) and retained (bottom) when a minimum threshold of 0.001 substitutions per site between divergence times is applied to trees randomly sampled from the prior distribution of the *M_IB_* model. Trees plotted using Gram (Version 4.0.0, Commit 02286362; Foster, 2018) and the P4 phylogenetic toolkit (Version 1.4, Commit d9c8d1b1; Foster, 2004).

### 2 Tables

Table S1. The data for all *Cyrtodactylus* samples included in our phylogenetic analyses are included in a tab-delimited text file available from the project repository and archived on Zenodo (Oaks and Wood, Jr., 2021): https://raw.githubusercontent.com/phyletica/gekgo/master/phycoeval-msg-assemblies/ipyrad-assemblies/sample-data/Cyrt_localities.tsv.

Table S2. The data for all *Gekko* samples included in our phylogenetic analyses are included in a tab-delimited text file available from the project repository and archived on Zenodo (Oaks and Wood, Jr., 2021): https://raw.githubusercontent.com/phyletica/gekgo/master/phycoeval-msg-assemblies/ipyrad-assemblies/sample-data/Gekko_localities.tsv.

**Table S3.** Summary of shared divergences in the maximum a *posteriori* (MAP) phylogeny estimated under the generalized tree model for *Cyrtodactylus* (Figure 6A) and *Gekko* (Figure 6C). See Figures S14 & S15 for the shared divergence labels.

| Shared divergence label | Number of nodes | Posterior prob. | Mean time | Time 95% HPDI |
| --- | --- | --- | --- | --- |
| C1 | 3 | 0.569 | 3.163 | 3.075–3.255 |
| C2 | 2 | 0.894 | 2.254 | 2.151–2.361 |
| C3 | 2 | 0.897 | 1.524 | 1.435–1.610 |
| C4 | 2 | 0.449 | 1.195 | 1.112–1.281 |
| C5 | 3 | 0.303 | 0.957 | 0.878–1.041 |
| G1 | 2 | 0.812 | 10.012 | 9.634–10.409 |
| G2 | 2 | 0.354 | 1.514 | 1.398–1.621 |
| G3 | 3 | 0.192 | 1.228 | 1.123–1.343 |
| G4 | 2 | 0.552 | 0.844 | 0.725–0.978 |
| G5 | 3 | 0.283 | 0.496 | 0.409–0.588 |

**Table S4.** Convergence statistics we calculated with `sumphycoeval` from MCMC samples collected with `phycoeval`. We ran each MCMC chain for 15,000 generations, sampling every 10 generations.

| Summary | *Cyrtodactylus* | *Gekko* |
| --- | --- | --- |
| Number of chains | 25 | 25 |
| Samples skipped per chain | 101 | 101 |
| Samples retained per chain | 1400 | 1400 |
| Total samples | 35000 | 35000 |
| ASDSF | 0.0027 | 0.00093 |
| Root age PSRF | 1.0000057 | 1.0001458 |
| Root age ESS | 33671.16 | 26844.22 |
| Population size PSRF | 1.0006948 | 1.0006295 |
| Population size ESS | 17542.23 | 13789.14 |

**Table S5.** Settings used for assembling RADseq loci for *Cyrtodactylus* and *Gekko*.

| ipyrad setting | Value |
| --- | --- |
| assembly_method | denovo |
| datatype | rad |
| restriction_overhang | TATG, |
| max_low_qual_bases | 5 |
| phred_Qscore_offset | 33 |
| mindepth_statistical | 6 |
| mindepth_majrule | 6 |
| maxdepth | 10000 |
| clust_threshold | 0.85 |
| max_barcode_mismatch | 0 |
| filter_adapters | 1 |
| filter_min_trim_len | 35 |
| max_alleles_consens | 2 |
| max_Ns_consens | 0.05 |
| max_Hs_consens | 0.05 |
| min_samples_locus | 4 |
| max_SNPs_locus | 0.2 |
| max_Indels_locus | 8 |
| max_shared_Hs_locus | 0.5 |
| trim_reads | 0, 0, 0, 0 |
| trim_loci | 0, 0, 0, 0 |

### 3 The generalized tree model

Let *T* represent a rooted, potentially multifurcating tree topology with *N* tips and *n*(*t*) internal nodes ***t*** = *t*_1_,*t*_2_, … *t_n_*(*t*), where *n*(*t*) can range from 1 (the “comb” tree) to *N* – 1 (fully bifurcating, independent divergences). Each internal node *t* is assigned to one divergence time *τ*, which it may share with other internal nodes in the tree. We will use ***τ*** = *τ*_1_, …, *τ_n_*_(*τ*)_ to represent *n*(*τ*) divergence times, where *n*(*τ*) can also range from 1 to *N* – 1, and every *τ* has at least one node assigned to it, and every node maps to a divergence time more recent than its parent (Figure S17). Note, the number of divergence times is constrained by the number of internal nodes; *n*(*τ*) ≤ *n*(*t*). For convenience, we will index each *τ* from youngest to oldest. We assume the tree is ultrametric; all tips are at time zero, which we will denote as *τ*_0_.

**Figure S17.**
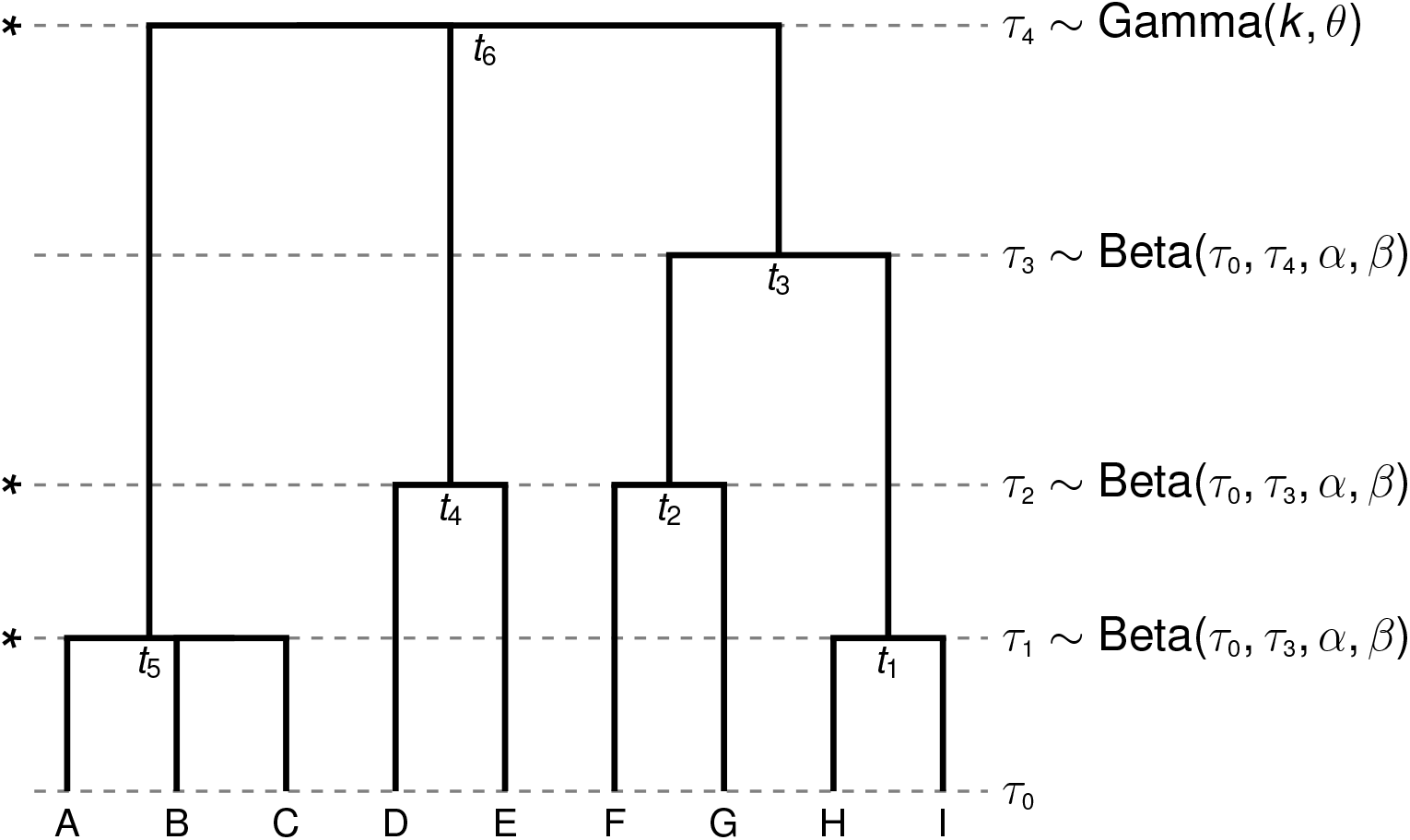
An illustration of the generalized tree model implemented in `ecoevolity`. The prior distributions of the divergence times are shown to the right, and “splittable” divergence times are indicated with an asterisk to the left. Figure created using Gram (Version 4.0.0, Commit 02286362; Foster, 2018) and the P4 phylogenetic toolkit (Version 1.4, Commit d9c8d1b1; Foster, 2004).

We assume all possible topologies (*T*) are equally probable; see Figure S1 for an example of the sample space of topologies under the generalized tree model. We assume the age of the root node follows a parametric distribution (e.g., a gamma distribution), and each of the other divergence times is beta-distributed between the present (*τ*_0_) and the age of the youngest parent of any node mapped to the divergence time. For example, in Figure S17, the parents of the nodes mapped to *τ*_1_ are *t*_3_ (parent of *t*_1_) and *t*_6_ (parent of *t*_5_), which are mapped to *τ*_3_ and *τ*_4_, respectively, the younger of which is *τ*_3_. Thus, divergence time *τ***_1_** in Figure S17 follows a beta distribution that is scaled to the interval between 0 and *τ*_3_, resulting in the prior probability density

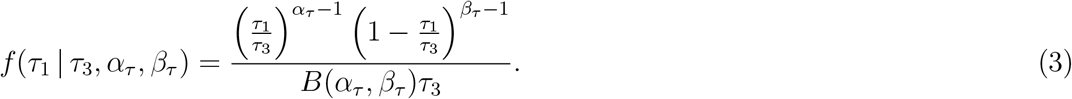

where *α_τ_* and *β_τ_* are the two positive shape parameters of the beta prior probability distribution, and *B*(*α_τ_, β_τ_*) is the beta function that serves as a normalizing constant. If we use *y*(*τ_i_*) to denote the divergence time of the youngest parent of any node mapped to the non-root divergence time, *τ_i_*, we can generalize this beta prior probability density as

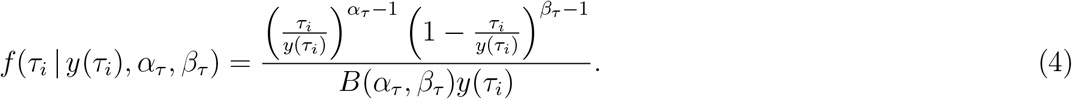

For additional flexibility, we allow a distribution to be placed on the alpha parameter of the beta distributions on the non-root divergence times (*α_τ_*). However, we constrain *β_τ_* = 1, simplifying the prior probability density of each non-root divergence time to

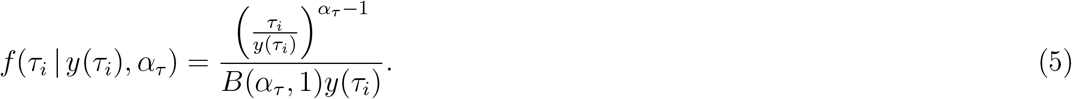

For all of our analyses presented in this paper, we constrained *α_τ_* = 1, which further simplifies the prior probability of each non-root divergence time to the uniform density

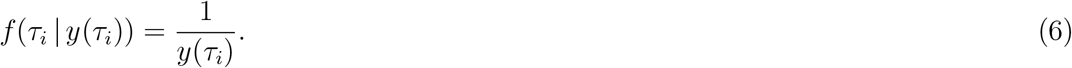

For simplicity below, we denote the probability density of a divergence time as *f*(*τ_i_* | *y*(*τ_i_*)), but our work generalizes to any proper probability distributions on the divergence times, including the full beta distribution (Equation 4).

### 4 Approximating the posterior of the generalized tree model

We use Markov chain Monte Carlo (MCMC), specifically Metropolis-Hastings (MH) (Metropolis et al., 1953; Hastings, 1970) algorithms, to sample from the generalized tree distribution. An MH algorithm works by stochastically proposing changes to parameters of a model, and using the following rule to determine the probability of accepting these moves:

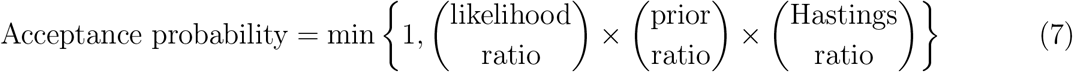

The product of the first two terms (likelihood and prior ratios) gives us the posterior density of the state of the model being proposed divided by the posterior density of the current state of the model. The Hastings ratio corrects for asymmetry in the proposal distribution by dividing the probability density of proposing the move that would reverse the move being proposed by the probability density of the move being proposed.

Below we describe MH moves for sampling from the generalized tree distribution, and derive the prior and Hastings ratio for each. These moves can be coupled with any likelihood function to calculate the likelihood ratio and sample from the posterior distribution of generalized trees using MH (Equation 7). In Sections 4.1–4.4, we describe a pair of reversible-jump moves, “split-time” and “merge-times,” for moving between trees with different numbers of divergence times. These are the core moves that allow the full space of the generalized tree distribution to be traversed and sampled. In Section 4.5, we describe moves that propose changes to the topology, but do not change the number of divergence-time parameters, nor their values. In Section 4.6, we describe a move that updates the values of divergence times without changing the topology (though we do describe an extension of this move that does update the topology).

#### 4.1 Split-time move

To generalize the space of trees, we introduce two reversible-jump moves: “split-time” and “merge-times.” When a reversible-jump move is to be attempted, the merge-times or split-time move is chosen with probability 0.5, except for two special cases:

1. If the current state of the chain is the most general tree model (*n*(*τ*) = *N* – 1), then the merge-times move is chosen with probability 1.
2. If the current state of the chain is the “comb” tree (*n*(*τ*) = 1), then the split-time move is chosen with probability 1.

The basic idea is to randomly divide a “splittable” divergence time into two nonempty sets of nodes, and assign one of the sets to a new, more recent divergence time. A divergence time is considered splittable if it has (1) more than one node mapped to it, or (2) a single multifurcating node mapped to it. For example, in Figure S17, divergence times *τ*_1_, *τ*_2_, and *τ*_4_ are splittable. The first step is to randomly choose a divergence time, *τ_i_*, from among the splittable divergence times. After dividing *τ_i_* into two sets of nodes, we need to randomly select a time more recent than *τ_i_* to assign one of the sets.

##### 4.1.1 Drawing the new divergence time

To get the new, proposed divergence time, we randomly draw a new divergence time between *τ*_*i*–1_ and *τ_i_* from a proposal distribution, where

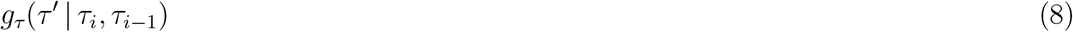

is the conditional probability density of proposing the new time *τ*′ given the times from the current values of *τ_i_* and *τ*_*i*–1_. In our implementation, we use a beta probability distribution scaled and shifted to the interval *τ*_*i*–1_–*τ_i_* so that the probability density of the new time is

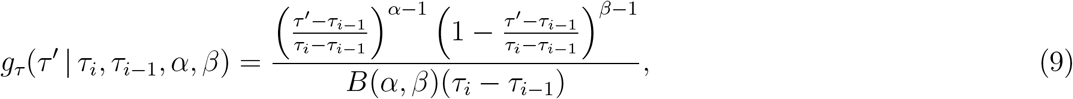

where *α* and *β* are the two positive shape parameters of the beta distribution, and *B*(*α,β*) is the beta function. For generality and simplicity, we will use *g_τ_* to denote this probability density of the proposed divergence time below.

##### 4.1.2 Prior ratio

The prior ratio for the split-time move is

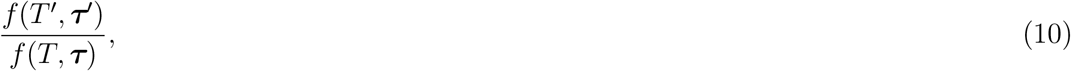

where *f*(*T*′, ***τ′***) is the prior probability of the proposed tree topology and divergence times. In our implementation, we assume that all possible tree topologies (across *n*(*τ*) = 1, 2, …, *N* – 1) are equally probable *a priori*. We also assume the divergence time of the root (*τ*_*n*(*τ*)_) is gamma-distributed, and each of the other divergence times is beta-distributed between the present (*τ*_0_) and the divergence time of the youngest parent of a node mapped to *τ_i_*, which we denote as *y*(*τ_i_*) (Figure S17; Equation 4). Given these assumptions, the prior ratio becomes

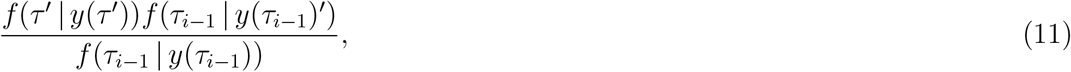

If *τ_i_* = 1 (i.e., the divergence time selected to split was the most recent divergence), then *τ*_*i*–1_ is the present, and so *f*(*τ*_*i*–1_ | *y*(*τ*_*i*–1_)′) = *f*(*τ*_*i*–1_ | *y*(*τ*_*i*–1_)) = 1. Also, if none of the nodes assigned to *τ*_*i*–1_ has a parent assigned to the newly proposed divergence time (i.e., *y*(*τ*_*i*–1_)′ ≠ *τ*′), then *y*(*τ*_*i*–1_)′ = *y*(*τ*_*i*–1_); e.g., in Figure S17, if *τ*_2_ is split, the prior probability density of *τ*_1_ is not affected, because the youngest parent of a node mapped to *τ*_1_ is mapped to *τ*_3_. In both of these special cases, the prior ratio further simplifies to

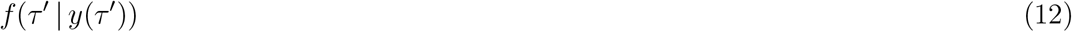

##### 4.1.3 Hastings ratio

The probability of proposing a split-time move involves several components. First, we have to choose to split rather than merge. We will account for the probability of this toward the end of this section. Next, we randomly choose a splittable divergence time *τ_i_* with probability 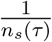, where *n_s_*(*τ*) is the number of splittable divergence times. As described in Section 4.1.1 above, we randomly choose a new divergence time *τ*′ more recent than *τ_i_* with probability density *g_τ_*.

When we divide *τ_i_* into two sets of nodes, if any polytomies get broken up, new branches will get added to the tree. Under certain models, each of these new branches will need values randomly drawn for parameters. For example, if using a “relaxed-clock” model, each new branch will need a substitution rate. Or, if using a multi-species coalescent model where each branch has its own effective population size, a value for this will need to be drawn. Note, this does not involve divergence-time parameters, because all nodes split from *τ_i_* will be assigned to *τ*′. We will use ***g**_z_* to represent the product of all the probability densities of the proposed values for the new branches. If no polytomies get broken up, or new branches created from broken polytomies do not require parameter values, then ***g**_z_* = 1.

We will deal with how the nodes assigned to *τ_i_* are divided into two sets below. For now, we will use Ξ to represent the probability of the proposed division of *τ_i_*. The probability density of the proposed split move is then

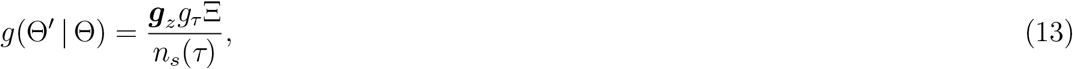

where Θ and Θ′ represent the full state of the model before and after the proposed split move, respectively. The move that would exactly reverse this split move would simply entail randomly selecting the proposed divergence time from all divergence times except the root, which would then be deterministically merged with the next older divergence time. The probability of this reverse move is

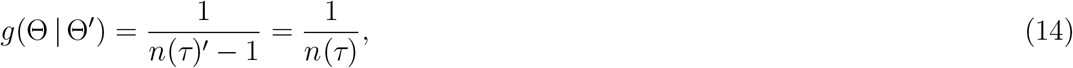

where *n*(*τ*) and *n*(*τ*)′ is the number of divergence times before and after the proposed split move, respectively.

The Hastings ratio for the split move is then

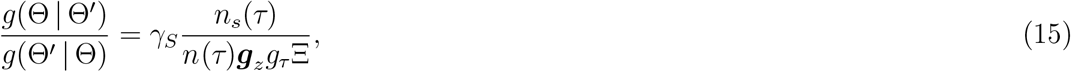

where *γ_S_* represents the probability of choosing to merge in the reverse move divided by the probability of choosing to split in the forward move, which is

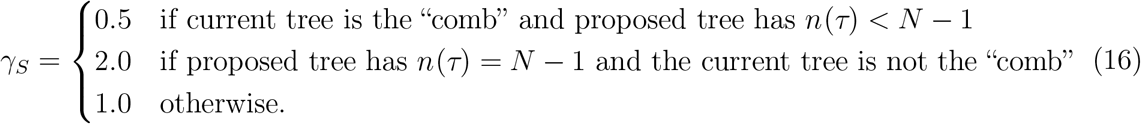

When we are working with a tree with more than three tips, the first case occurs whenever the current tree is the comb tree (*n*(*τ*) = 1), and the second case occurs whenever the proposed tree has no shared divergences nor multifurcations (*n*(*τ*) = *N* – 1). However, for full generality we need to include the second condition in both of these cases to account for the situation where we are working with a tree with only three tips.

#### 4.2 Merge-times move

In the “merge-times” move, we randomly choose *τ_x_* from one of the *n*(*τ*) – 1 non-root divergence times. Then, we merge *τ_x_* with the next older divergence time, *τ*_*x*+1_. This will create shared divergence times among nodes and/or multifurcating nodes. We will use 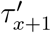 to refer to the newly merged divergence time proposed by the move.

##### 4.2.1 Prior ratio

Generally, the prior ratio for the merge-times move is the same as Equation 10. Assuming (1) all topologies are equally probable, (2) the divergence time of the root (*τ_n_*_(*τ*)_) is gamma-distributed, and (3) each of the other divergence times is beta-distributed between the present (*τ*_0_) and the age of the youngest parent to the nodes mapped to the divergence time (Figure S17; Equation 4), the prior ratio becomes

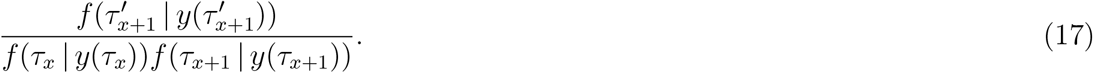

If *τ*_*x*+1_ is the root of the tree, then 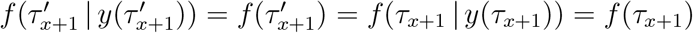, and this probability density is given by the gamma prior distribution on the divergence time of the root (Figure S17).

##### 4.2.2 Hastings ratio

The probability of the forward merge-times move is simply

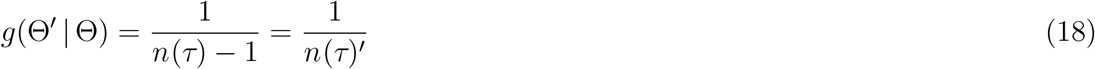

where *n*(*τ*) and *n*(*τ*)′ is the number of divergence times before and after the proposed merge move, respectively.

Borrowing from Equation 13, the probability density of the split move that would exactly reverse the proposed merge move is

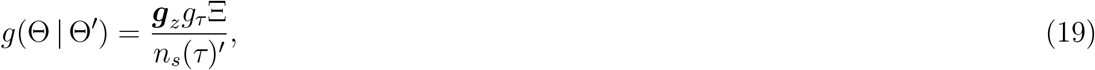

where *n_s_*(*τ*)′ is the number of splittable divergence times *after* the proposed merge-times move.

The Hastings ratio for the merge move is then

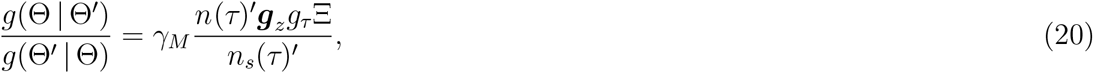

where *γ_M_* represents the probability of choosing to split in the reverse move divided by the probability of choosing to merge in the forward move, which is

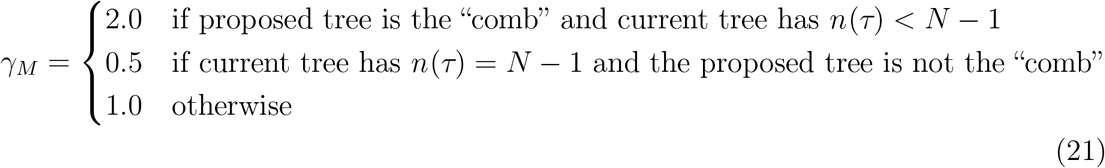

#### 4.3 Expanding Ξ

Up to this point, we have not dealt with how, during the split-time move, we divide the nodes mapped to *τ_i_* into two sets, one of which gets assigned to the new divergence time drawn between *τ*_*i*–1_ and *τ_i_*. This has to be done with care to ensure that every possible configuration of two divergence times derived from the nodes assigned to *τ_i_* can be proposed, such that it properly balances the reverse merge-times move. As above, we use Ξ to represent the probability of the proposed division of *τ_i_*’s nodes.

In the next two sections, we show how this is done for two special cases. The first special case illustrates how we first choose which nodes currently mapped to *τ_i_* will get moved to the new divergence time. The second special case shows how we handle any multifurcating nodes that have been chosen to be moved to the new divergence time. In the third section, we build on these special cases to show a general solution for Ξ.

##### 4.3.1 The case of all bifurcating nodes mapped to *τ_i_*

We will use *n*(*t* ↦ *τ_i_*) to represent the number of nodes mapped to *τ_i_*. If all *n*(*t* ↦ *τ_i_*) nodes mapped to *τ_i_* are bifurcating, we randomly divide these nodes into two non-empty sets and then randomly choose one of the two sets of nodes to move to the new divergence time. For example, this would be the case if *τ*_2_ is chosen to split from the tree shown in Figure S17.

The number of ways *n*(*t* ↦ *τ_i_*) can be divided into two non-empty subsets is given by the Stirling number of the second kind, which we denote as *S*_2_(*n*(*t* ↦ *τ_i_*), 2). We uniformly choose among these, such that the probability of randomly selecting any set partition of the *n*(*t* ↦ *τ_i_*) nodes mapped to *τ_i_* is 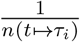. After partitioning the nodes into two sets, there is a 1/2 probability of choosing one set to move to the new, more recent divergence time. Thus, *when all of the nodes mapped to τ_i_ are bifurcating* the probability of each possible splitting of *τ_i_* is

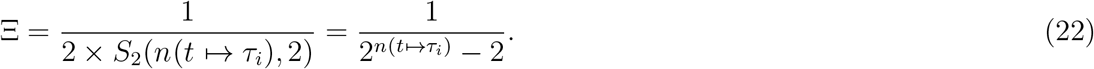

##### 4.3.2 The case of a single polytomy mapped to *τ_i_*

Next, let’s consider another special case where the number of nodes mapped to *τ_i_* is one (i.e., a single polytomy). For example, this would be the case if *τ*_4_ is chosen to split from the tree shown in Figure S17. In this case, we randomly resolve the polytomy, by randomly (uniformly) choosing a set partition of the descending branches into non-empty subsets. Any subsets with only one branch remain attached to the original polytomy node, while each subset with multiple branches get split off to form a new node (clade) that descends from the original polytomy node. These new nodes are assigned to the new, more recent divergence time *τ*′. The number of ways to partition the descending branches of the polytomy are thus *B_b_* – 2, where *B_b_* is the Bell number (Bell, 1934)—the number of possible set partitions of the *b* branches descending from the polytomy. We have to subtract 2 from *B_b_*, because we do not allow the two “extreme” set partitions with one or b subsets. The former would move the whole polytomy to the new divergence time, and the latter would leave the polytomy as is. Neither of these scenarios adds a dimension (divergence time) to the model. We avoid these two scenarios using rejection so that the remaining partitions of the b descending branches are chosen uniformly. Thus, *when only a single polytomy is mapped to τ_i_* the probability of each possible splitting of *τ_i_* is

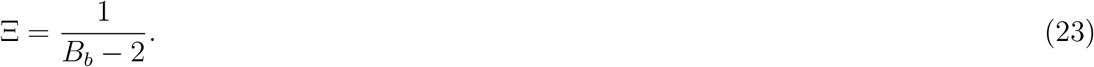

##### 4.3.3 The case when multiple nodes, including at least one polytomy, are mapped to *τ_i_*

When multiple nodes are mapped to *τ_i_*, and at least one is a polytomy, we need to do some more accounting to ensure that we can reach every possible arrangement of two divergence times that can be merged to form the current configuration of nodes mapped to *τ_i_*. Similar to the case with all bifurcating nodes, we will first divide the *n*(*t* ↦ *τ_i_*) nodes mapped to *τ_i_* into two subsets and randomly choose one of these subsets to move to the proposed, more recent divergence time, *τ*′. For each multifurcating node that ends up in the subset to be moved to *τ*′ (if any), we need to either break up the polytomy, as we did in the case of the single-polytomy case above, or move the entire polytomy to *τ*′.

Unlike in the case of only bifurcating nodes mapped to *τ_i_*, when we partition the *n*(*t* ↦ *τ_i_*) nodes mapped to *τ_i_* into two sets, we must allow for the case where all *n*(*t* ↦ *τ_i_*) end up in the set to move to *τ*. This is because, if any of the polytomy nodes get broken up, they will leave at least one node at *τ_i_*, and the dimension of the model will change (i.e., the number of divergence times will increase by one). So, we have to allow an empty subset when we randomly partition the *n*(*t* ↦ *τ_i_*) into two subsets. However, we cannot allow the empty subset to be chosen to move to *τ*′. There are *S*_2_(*n*(*t* ↦ *τ_i_*), 2) + 1 ways to partition the *n*(*t* ↦ *τ_i_*) nodes mapped to *τ_i_* into two subsets if we allow the set partition with one empty subset. For each of these, there are two ways to choose the subset to move to *τ*′, and of all of these, there is one scenario we will reject: if the empty set gets selected to move to *τ*′. Thus, there are

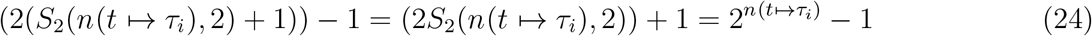

ways to choose a subset of the nodes assigned to *τ_i_* for moving to the new divergence time, and the probability of each is

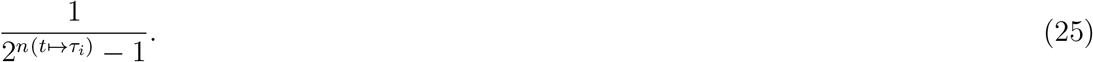

For each polytomy mapped to *τ_i_* that ends up in the set of nodes to move to *τ*′ (if any), we randomly choose one of the *B_b_* possible set partitions of the *b* branches descending from the polytomy. However, we will reject the set partition with *b* subsets (i.e., all branches end up in their own subset). We reject this, because no subclades get broken off from the polytomy to move to *τ*′, and this scenario is already taken into account by the polytomy node not ending up in the set of nodes to move to *τ*′ in the first place. However, we need to allow the scenario where all *b* branches descending from a polytomy get assigned to a single set, which results in the entire polytomy node getting moved to *τ*′, as long as at least one node remains assigned to *τ_i_* (we will handle this in a bit). Thus, for each polytomy in the set of nodes to be moved to *τ*′, there are *B_b_* – 1 ways to move it. Using *n_p_*(*t* ⟹ *τ*′) to represent the number of polytomies in the subset of nodes to be moved to *τ*′, the total number of ways these polytomies can be moved to *τ*′ is

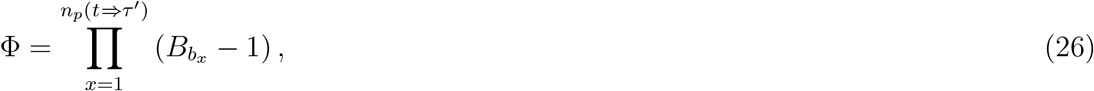

and the probability of each is equal to

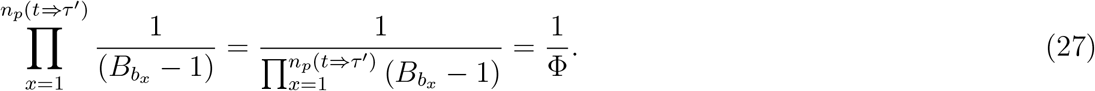

If no polytomies end up in the subset of nodes to move to *τ*′, then Φ = 1.

However, if all *n*(*t* ↦ *τ_i_*) nodes mapped to *τ_i_* end up in the set of nodes to be moved to *τ*′, we need to reject the case where none of the polytomy nodes gets broken up (i.e., for every polytomy, all the descending branches get partitioned into a single set), because no nodes would remain assigned to *τ_i_*, and the move would simplify to changing the value of *τ_i_*. Thus, if all *n*(*t* ↦ *τ_i_*) nodes mapped to *τ_i_* end up in the set of nodes to move to *τ*′, the total number of ways all *n_p_*(*t* ⇒ *τ*′) polytomies can be moved to *τ*′ is

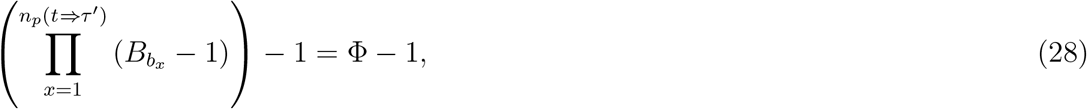

and the probability of each is equal to

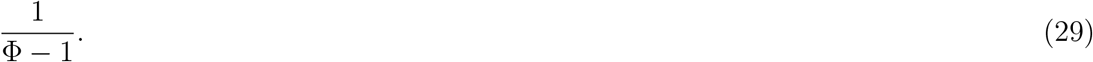

Given all of this, the probability of choosing a subset of nodes from *τ_i_* to move to the new divergence time across all possible cases is

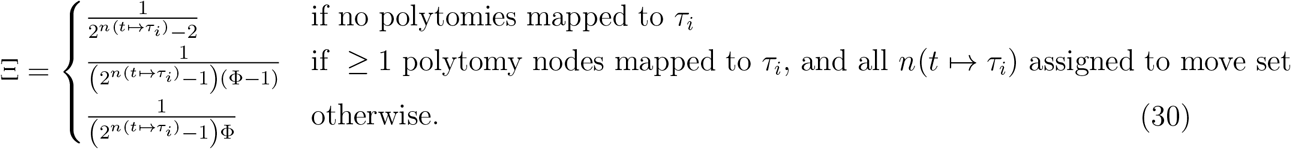

Notice, the case of *n*(*t* ↦ *τ_i_*) = 1 (i.e., only a single polytomy node mapped to *τ_i_*) is simply a special case of the second condition above, where all the nodes assigned to *τ_i_* end up in the set to move to the new divergence time, including at least one polytomy.

#### 4.4 Validation of Split-time and Merge-times moves

To validate the split-time/merge-times moves, we used them to sample from the prior distribution of trees with 5, 6, and 7 leaves. If working correctly, we should sample all *n*(*T*) tree topologies with an equal frequency of 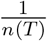. If we collect 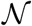 MCMC samples from the prior distribution, the number of times a topology is sampled should be approximately distributed as Binomial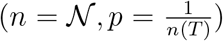; i.e., binomially distributed where the number of “trials” is equal to the number of samples, and the probability of sampling each topology is 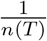. We found a close match between the number or times each tree was sampled by our reversible-jump MCMC chain and the expected number, and failed to reject the expected binomial distribution using χ^2^ goodness-of-fit test (Figure S18; *p* = 0.742, 0.464, and 0.172 for the test with a 5, 6, and 7-leaved tree).

**Figure S18.**
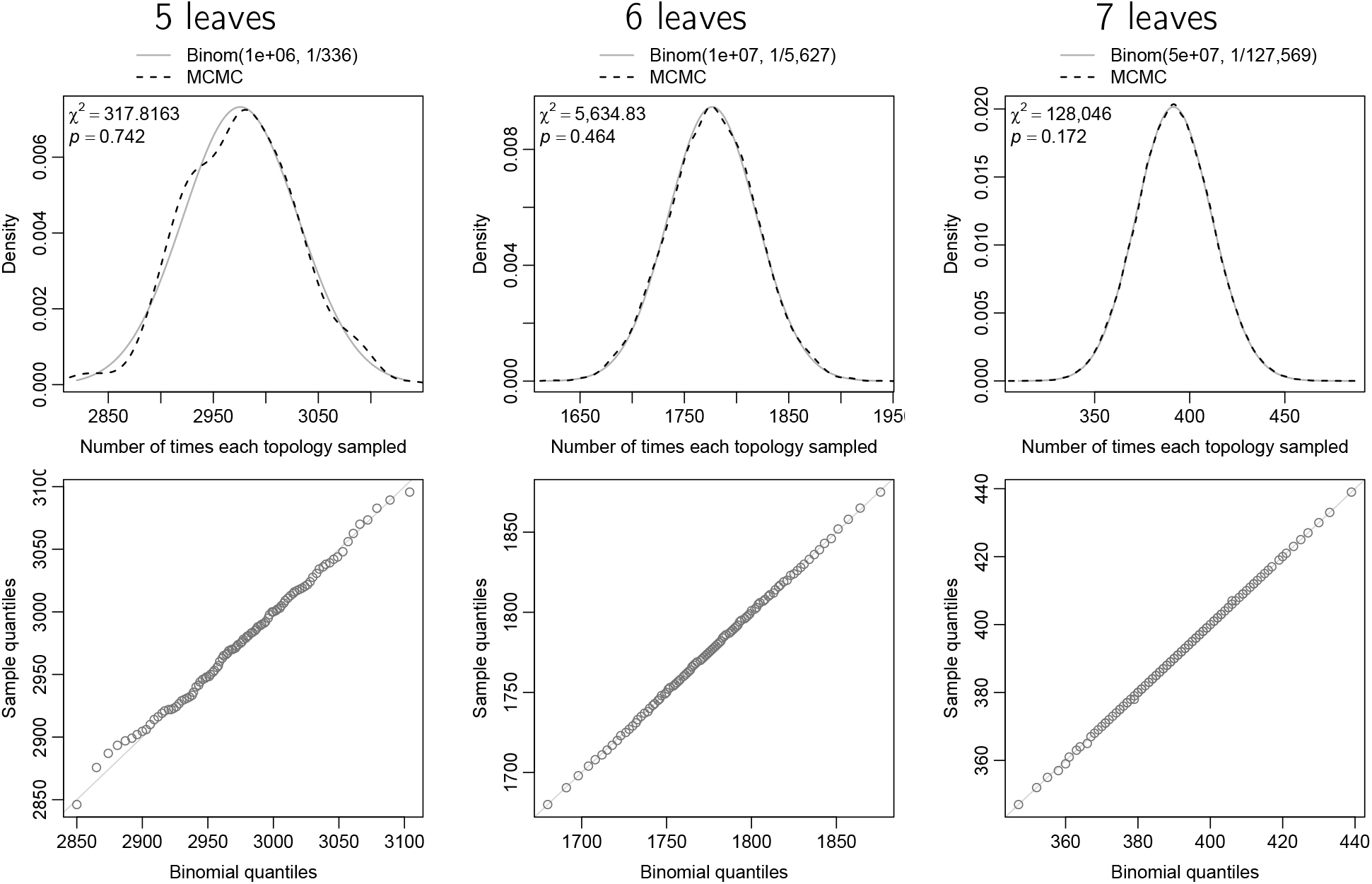
Comparing the expected to the observed number of times each topology is sampled by an MCMC chain using our split-time and merge-times moves. Under our generalized tree distribution, how often each topology is sampled should follow a 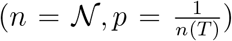 distribution, where 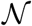 is the total number of MCMC samples.

#### 4.5 Nested-neighbor-node-swap move

The split-time and merge-times moves can sample all of the space of the generalized tree distribution (assuming the age of the root node is fixed). However, we implemented additional topology moves that do not jump between tree models with different numbers of divergence times.

The goal of this move is to change the topology without changing the number or timing of divergences. We start by randomly picking a non-root divergence time, *τ_i_*, with probability 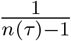. Next, we find the divergence time, *τ_j_*, that contains the node that is the youngest parent of nodes mapped to *τ_i_*. We then randomly pick one of the nodes mapped to *τ_j_* that has children mapped to *τ_i_*, we will call it *t_a_*. Each child of *t_a_* that is mapped to *τ_i_* will randomly contribute one of its children to a “swap pool” of nodes. If *t_a_* has children that are *not* mapped to *τ_i_*, we randomly pick one of these children and add it to the swap pool *if* it is younger than *τ_i_*. If the selected child of *t_a_* is older than *τ_i_*, we randomly sample one of its children and continue to do so until we have chosen a descendant node that is younger than *τ_i_*, which we then add to the swap pool. Lastly, we randomly pick two nodes from the swap pool and we swap their parents.

After the proposed move, the structure of the tree rootward of the swapped nodes is the same. Because of this, the move that would reverse the proposed move would be equally probable; it would involve (1) choosing the same non-root divergence time *τ_i_*, (2) choosing the same *t_a_*, (3) choosing the children that swapped parents in the forward move to enter the swap pool, and (4) picking the nodes that swapped parents in the forward move from the pool and swapping their parents back. In #3, all the parent nodes involved have the same number of children as before the proposed forward move, and so the probability of the reverse move will be equal. As a result, the Hastings ratio for the move is 1.

For example, for the tree in Figure S17 if we randomly selected *τ*_1_, the divergence containing the youngest parent of the nodes mapped to *τ*_1_ is *τ*_3_. Divergence time *τ*_3_ only has one node that is a parent of nodes assigned to *τ*_1_, which is *t*_3_. Node *t*_3_ only has one child mapped to *τ*_1_ (*t*_1_), which will randomly contribute one of its children, Leaf H or I, to the swap pool. Node *t*_3_ also has children that are not mapped to *τ*_1_, one child *t*_2_ that is mapped to *τ*_2_. Node *t*_2_ is considered for the swap pool, but it is too old (it is older than *τ*_1_ and thus could not become a child of *t*_1_. So, we randomly consider one the children of *t*_2_, Leaf F or G, for the swap pool, either of which is young enough to be added to the swap pool. Next, we randomly choose two nodes from the swap pool, which has exactly two nodes in this case, a child of *t*_1_ (Leaf H or I) and *t*_2_ (Leaf F or G), and these nodes swap parents. If we assume that Leaves G and H were swapped, it is clear that the probability of the reverse move that would swap them back is equally probable.

We also implemented variations of this move that make larger changes to the tree topology. For example, we can perform the swap for all of the nodes mapped to *τ_j_* that have children mapped to *τ_i_* (instead of randomly choosing one of them). Another option is to randomly permute the parents of all of the nodes in the swap pool, rather than swap the parents of just two of the nodes. By chance, when doing this permutation of the nodes in the swap pool, it is possible to end up with the same topology we started with. To avoid proposing the same state, we iteratively permute the parents of the nodes in the swap pool until we have a new topology (the parents of at least some of the nodes in the swap pool have changed). Just like with the swap move, this permutation move can be performed on one randomly selected node of *τ_j_* that has children mapped to *τ_i_*, or to all of them.

##### 4.5.1 Validation of move

To validate this move, we used it to sample from a uniform distribution over the topologies of a 6-leaved bifurcating tree. There are 945 topologies for a rooted, bifurcating tree (Felsenstein, 1978). If the move is working correctly, the number of times we sample each of them should follow a 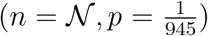 distribution, where 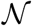 is the total number of MCMC samples. From an MCMC sample of 100,000 trees, we found a close match to this expected distribution, and were unable to reject it using a χ^2^ goodness-of-fit test (Figure S19; *p* = 0.51).

**Figure S19.**
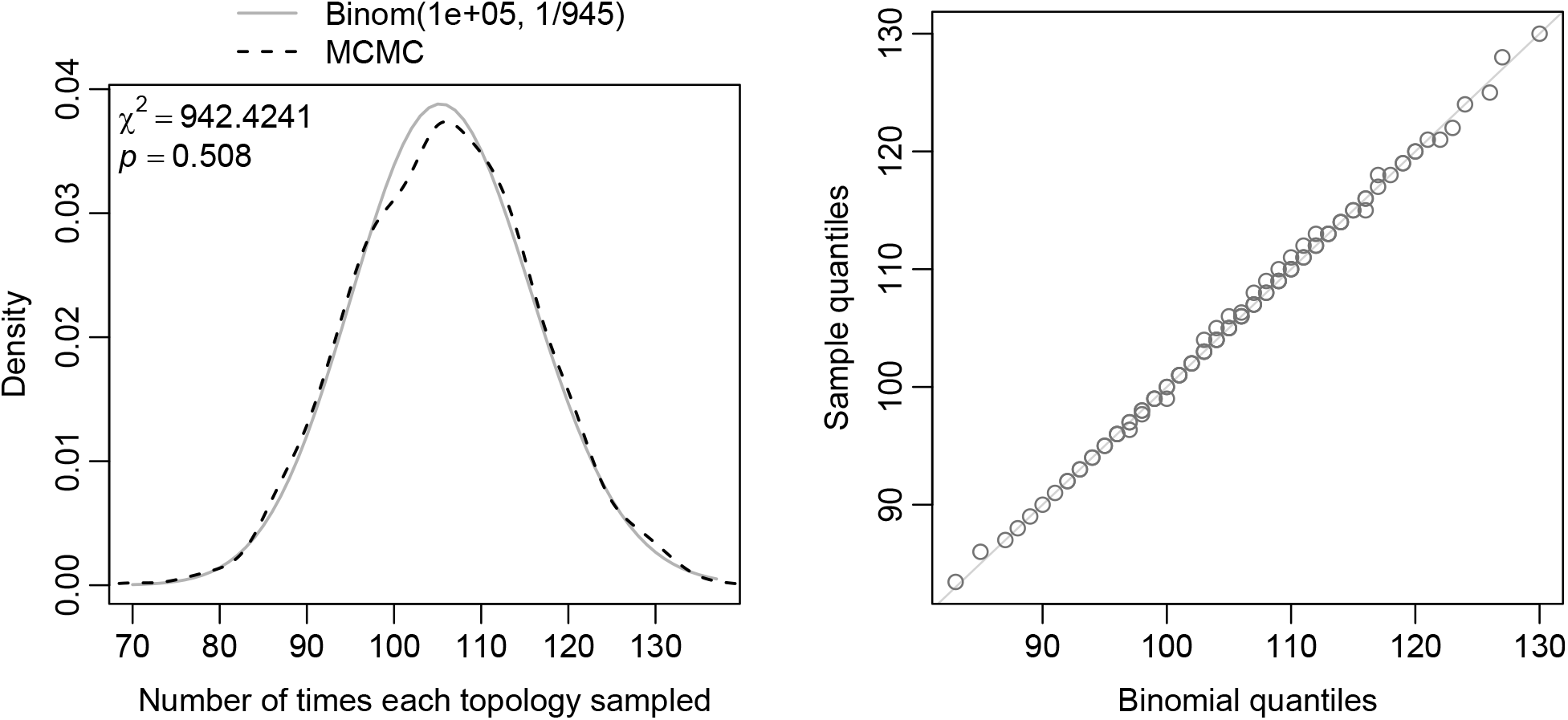
Comparing the expected to the observed number of times each topology of a rooted, 6-leaved, bifurcating tree is sampled by our nested-neighbor-node-swap move. Our MCMC sample of 100,000 trees closely matched the expected 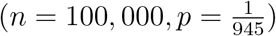 distribution.

#### 4.6 Divergence time slide bump move

To begin this move, we randomly pick one of the divergence times, *τ_i_*. Next, we draw a uniform deviate, *u* ~ Uniform(−λ, λ), where λ is a tuning parameter that can be adjusted to improve the acceptance rate of the proposal. Then, we get a new divergence time value by *τ_i_e^u^*. We will index our randomly selected divergence time, *τ_i_*, as *τ*_1_. We then use *τ*_1_, *τ*_2_, …, *τ_n_* to represent the selected time, *τ*_1_, and all the divergence times between *τ*_1_ and *τ*_1_*e^u^* that contain nodes ancestral or descendant to the nodes mapped to *τ*_1_. Note, that incrementing indices count younger or older divergence times, depending on whether *τ_i_e^u^* < *τ*_1_ or *τ_i_e^u^* > *τ*_1_, respectively.

The simplest case is that we do not have any intervening divergence times, and so we only have *τ*_1_. This will happen when *τ*_1_*e^u^* is older than the oldest node that is a child of the nodes mapped to *τ_i_* and younger than the youngest node that is parent of the nodes mapped to *τ_i_* In that case, we propose a new time to which to slide *τ*_1_ as

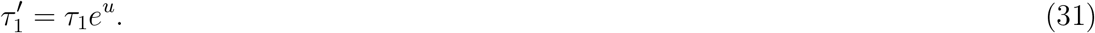

To reverse this move (slide *τ*_1_ back) would be

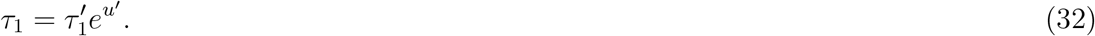

To solve for the uniform deviate that would exactly reverse the move (*u*′), we take the log of Equation 32 and solve for *u*′,

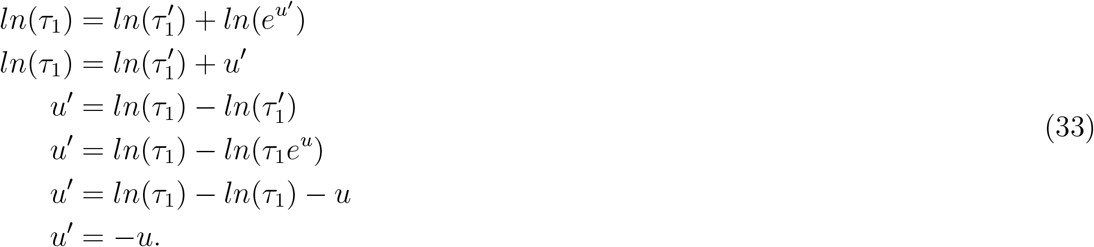

To get the Hastings ratio for this move, we use the formula of Green (1995),

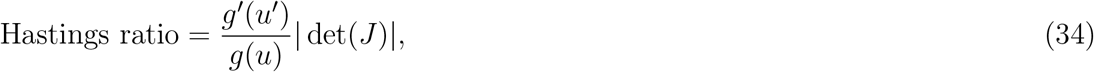

which is the ratio of the probability of drawing the random deviate that would reverse the proposed move to the probability of drawing the random deviate of the proposed move, multiplied by the absolute value of the determinant of a Jacobian matrix. Because the forward and reverse random deviates are uniform, 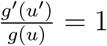, and the Hastings ratio reduces to just the Jacobian term,

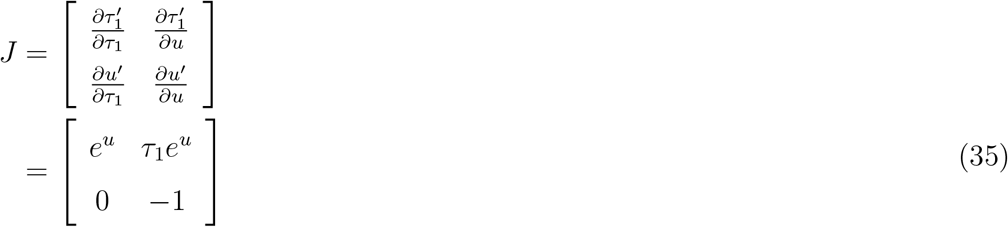

In the next simplest case, there is one intervening divergence time *τ*_2_. In this case, *τ*_1_ will slide to *τ*_2_ and “bump” it to the new time *τ*_1_*e^u^*. More formally, the move will be

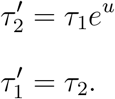

Again, the uniform deviate that would exactly reverse this move would be

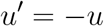

and the reverse move would be

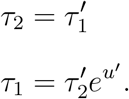

Again, the Hastings ratio reduces to just the Jacobian term,

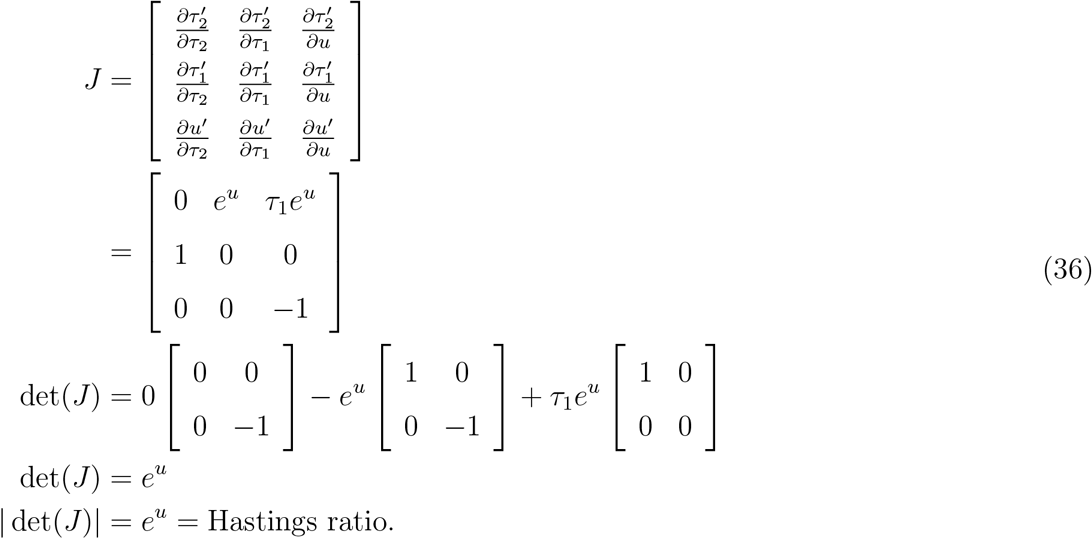

To generalize this to an arbitrary number of intervening divergence times that will be bumped, we have

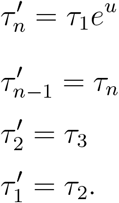

Again, the uniform deviate that would exactly reverse this move would be

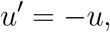

and the reverse move would be

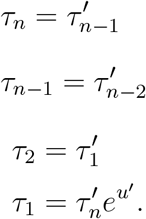

The Hastings ratio reduces to just the Jacobian term,

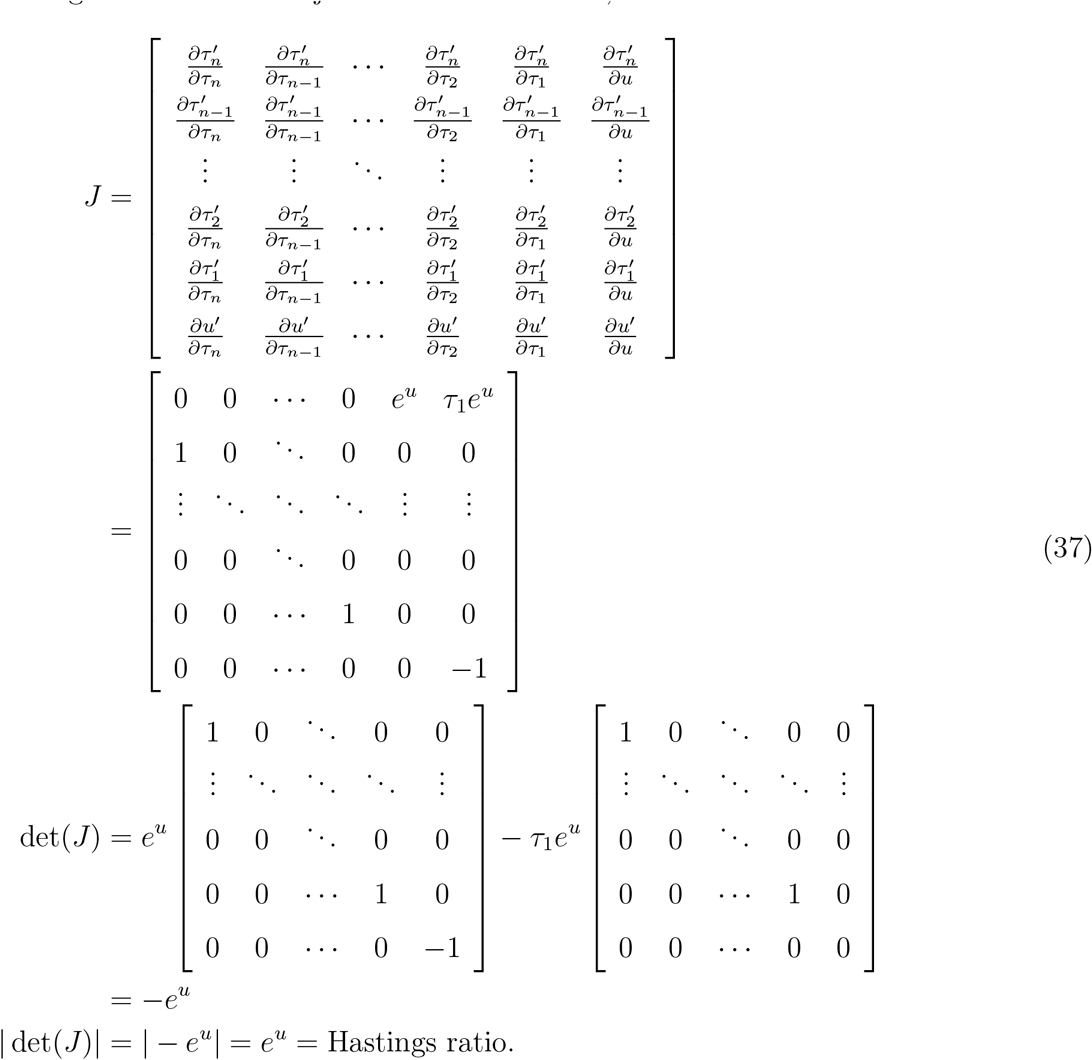

We avoid values of *τ*_1_*e^u^* that are less than zero, by rejecting the proposed move. There is no upper limit for the move, because the root of the tree can be moved to an arbitrarily old divergence time. However, in our implementation, the prior on the divergence time of the root is different than the other divergence times, and can be much more informative. In such cases, we might be able to improve mixing and tuning of the move be excluding the root divergence time from the move. We do this by selecting only non-root divergence times, and rejecting any proposed moves where *τ*_1_*e^u^* is older than the root.

##### 4.6.1 An extension to this move

We can easily extend this move to also propose new topologies. Whenever we have a “bump” that involves a node and its children, we can propose a node swapping or permuting move described above (see the nested-neighbor-node-swap move and its extensions). Because we are sliding the nodes to take the position of the nodes they bump, the swap or permute moves are simplified a bit. We do not have to worry about *τ_j_* contributing a child that is older than its potential new parents, so we never need to randomly choose descendants until we find a node that is younger than *τ_i_*. We implemented this move, but jointly proposing changes to continuous divergence time parameters and changing the topology might lead to poor acceptance rates (Yang, 2014), so using separate moves to update divergence times and the topology is likely a better strategy.

##### 4.6.2 Validation of this move

We used this move to sample from the prior distribution to ensure that the distribution of sampled divergence times matched the gamma-distributed prior we placed on the root age and the beta priors we placed on all other divergence times. To validate the extension of this move that also incorporates node swapping when nodes “bump,” we used it to sample from a uniform distribution over the topologies of a 6-leaved bifurcating tree. If the move is working correctly, the number of times we sample each of the 945 topologies should be follow a 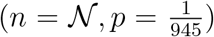 distribution, where 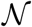 is the total number of MCMC samples. We found a close match between our samples and this expected distribution, and could not reject it using a χ^2^ goodness-of-fit test (Figure S20; *p* = 0.165,).

**Figure S20.**
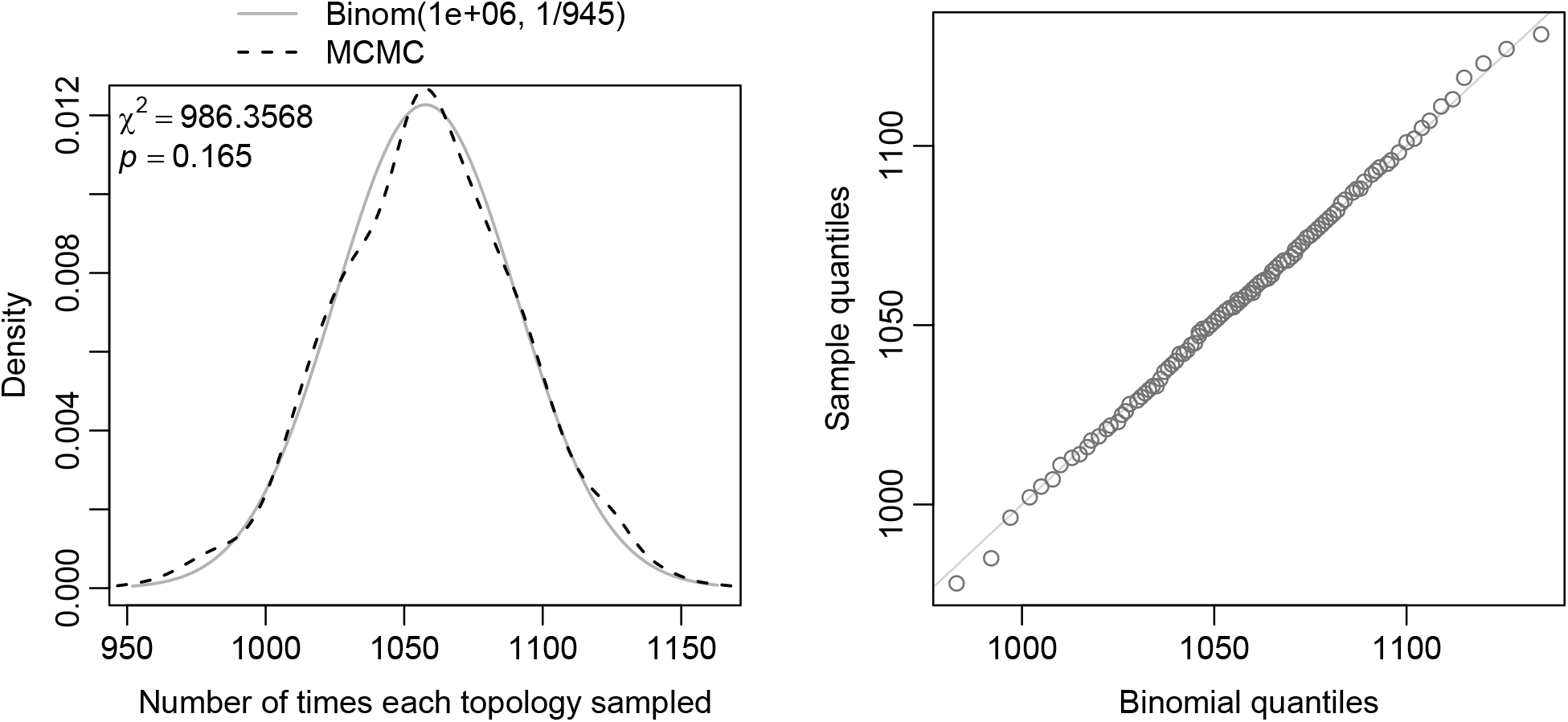
Comparing the expected to the observed number of times each topology of a rooted, 6-leaved, bifurcating tree is sampled by the node-swapping extension to our divergencetime-slide-bump move. Our MCMC sample of 1 million trees closely matched the expected 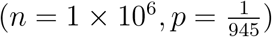 distribution.

